# Loratadine decreases methicillin-resistant *Staphylococcus aureus* virulence through global hemolysin regulators *vraSR* and *agr*

**DOI:** 10.1101/2025.06.05.657702

**Authors:** Chase Dillon, Isabel Marshall, Maggy Henkel, Halie Balogh, Robin Stempel, E. Renee Monge, Gabriel Z. Valenzano, Meghan S. Blackledge, Heather B. Miller

## Abstract

Methicillin-resistant *Staphylococcus aureus* (MRSA) is a bacterial pathogen that relies on expression of a wide range of virulence factors. Hemolysins are a diverse collection of cytolytic toxins that promote virulence by being excreted and lysing host blood cells. We previously reported on novel anti-virulence compounds that modulate hemolysis in vitro, including a brominated carbazole (compound 8) and the active ingredient in Claritin (loratadine). We analyzed expression and activity of MRSA hemolysins in two hospital-acquired strains. Even with the same staphylococcal cassette chromosome *mec* (SCC*mec*) type, the strains displayed unique responses to loratadine where both repression and elevation of hemolysis occurred. These compounds are potentially binding and regulating the master regulatory protein, Stk1, which has known ties to hemolysis in *S. aureus*. We hypothesized that modulation of hemolysis by these putative Stk1 inhibitors would extend to community-acquired strains of MRSA. Furthermore, we wanted to determine if Stk1 was the only hemolysis regulatory protein being engaged by these compounds. To test this, we examined compound 8 and loratadine’s effects on two different USA300 strains, in addition to transposon mutants for a range of hemolysis regulatory genes. We observed strain-specific regulation of hemolysis by Stk1. Loratadine continued to downregulate both alpha and beta hemolytic activity in USA300 JE2 in both Stk1-dependent and independent pathways. Additionally, we report that this anti-virulence compound effectively disrupts hemolysis by modulating both *agr* and *vraSR* global regulators, contributing to widespread cytotoxin repression. Both alpha hemolysin (*hla)* and RNAIII/delta hemolysin *(hld)* mRNA reductions occurred in vitro and in a human cell line model of MRSA infection. Together, these results expand our knowledge of loratadine’s mode of action as a virulence modulator.

## Introduction

*Staphylococcus aureus* is a common pathogen of humans and livestock causing infections of the skin, soft tissue,^1^ blood,^2^ and lungs,^3^ among other complications. To optimize colonization and survival in the host, *S. aureus* encodes a vast array of virulence factors. Among the most detrimental virulence factors are hemolysins, which are cytolytic toxins that target and damage red blood cells. Hemolysis allows the bacteria to scavenge heme from host cells, aiding in bacterial proliferation and spread of infection.^4^ The four hemolysins in *S. aureus* are primarily secreted during post-exponential phase growth.^5–7^ Each hemolysin has a different mechanism of action, but they collectively contribute to virulence. Alpha hemolysin (alpha toxin) is encoded by *hla* and beta hemolysin (a sphingomyelinase also known as phospholipase C) is encoded by *hlb.*^8, 9^ Gamma hemolysins have three components, A, B, and C, encoded by *hlgA*, *hlgB*, and *hlgC*, respectively.^10^ Finally, delta hemolysin, encoded by *hld,* is a peptide and also considered a phenol-soluble modulin (PSM).^11^

Hemolysin levels are controlled by an intricate network of regulatory proteins. Global regulators of hemolysis include at least three systems. The *agr* (accessory gene regulator) locus consists of *agrBDCA*, which is transcribed from one unit known as RNAII. Additionally, the *agr* locus has an adjacent transcriptional unit transcribed in the opposite direction called RNAIII. Sharing RNAIII’s promoter, *hld* is encoded within RNAIII.^12, 13^ The *agr* locus is well-studied and regulates over 100 genes in *S. aureus*. In addition to its role in control of quorum sensing, it also regulates virulence genes by upregulating exotoxins such as alpha, beta, gamma, and delta hemolysins.^14, 15^ Secondly, the Sar (staphylococcal accessory gene regulator) protein family consists of 11 DNA-binding proteins homologous to SarA, which regulate virulence gene expression.^16–18^ SarA has long been known to upregulate the expression of all hemolysins in *S. aureus*^15^ as well as contribute to PSM levels.^19, 20^ Third, the two-component system (TCS) SaeRS (staphylococcal accessory protein effector) globally regulates hemolysis through the sensor histidine kinase, SaeS, that detects environmental signals and phosphorylates SaeR, the cognate response regulator, leading to transcription of target genes including *hla* and *hlb.* ^21–23^ The master regulatory kinase Stk1 (eukaryotic-like serine-threonine kinase 1) also plays an important role in *S. aureus* virulence regulation,^24, 25^ although its exact role in hemolysis has not been fully elucidated. Several groups have reported that deletion of *stk1* increased hemolysis.^26–28^ These results supported Stk1 suppressing these virulence factors. However, there is conflicting data regarding Stk1’s role, with other reports showing the protein promoting hemolysis.^29, 30^ Strain-specific differences may be the cause of these contradictions; alpha toxin in particular has been shown to be regulated in a strain-specific fashion.^20, 29^

Our group has previously reported investigations surrounding two putative Stk1 inhibitors. The first is a brominated carbazole (compound 8, Figure 2C)^31^ and the second is a structurally related tricyclic molecule, loratadine, the active ingredient in Claritin (Figure 2D).^32^ Both compound 8 and loratadine have shown effective beta-lactam antibiotic potentiation in multiple strains of methicillin-resistant *S. aureus* (MRSA).^31, 32^ Furthermore, loratadine has been shown to prevent biofilm formation in vitro.^32, 33^ Combined with genetic results comparing wild type MRSA strains to *stk1* deletion mutants, we concluded that Stk1, with known ties to both antibiotic resistance and biofilm formation, was the likely target of compound 8^31^ and loratadine.^32^ We demonstrated that the *mec* and *bla* operons were both being modulated with these antibiotic adjuvants, leading to β-lactam potentiation.^31, 32^

**Figure 1:**
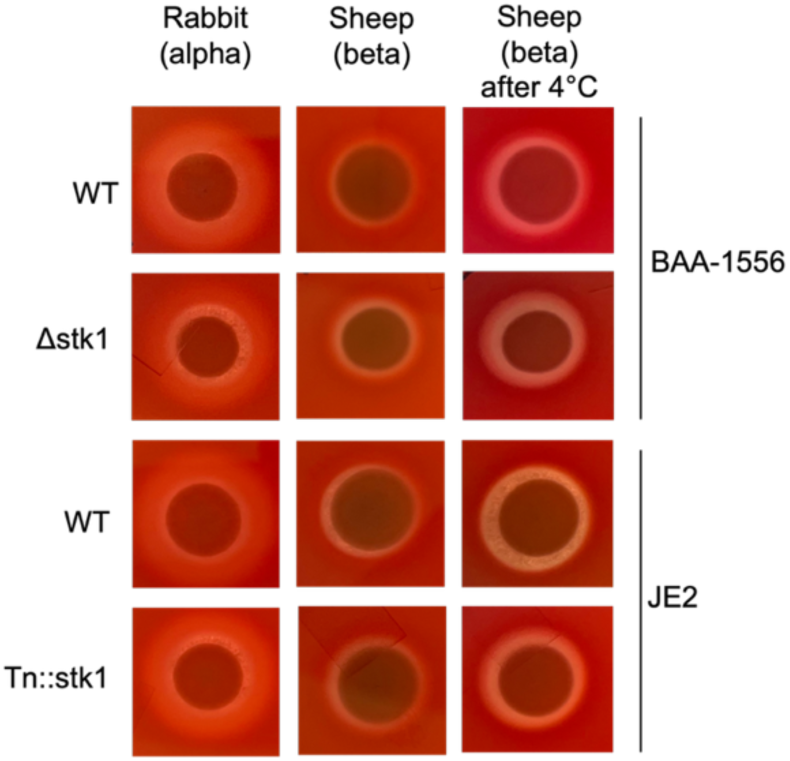
Blood agar reveals Stk1-mediated hemolysis regulation that is strain-specific. Each strain was grown on either rabbit blood or sheep blood agar. Only sheep blood agar plates were imaged again after 4 °C incubation overnight.

**Figure 2:**
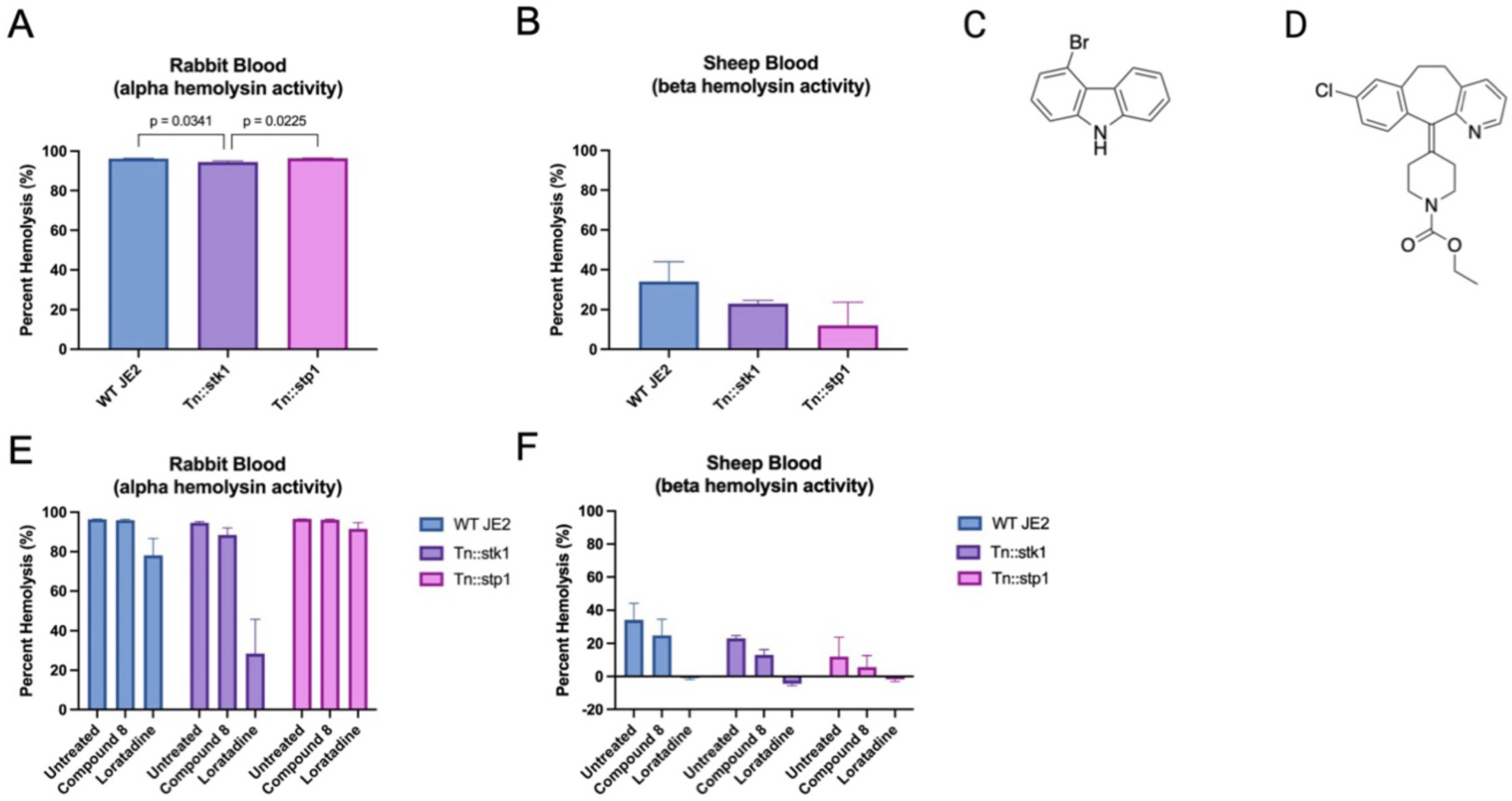
Quantitative hemolysis assays using (A) rabbit or (B) sheep blood reveal Stk1 promoting alpha hemolytic activity in USA300 strain JE2. One-way ANOVAs were used to assess variance between means of every sample. Only adjusted p values ≤0.05 are displayed. (C) Chemical structure of compound 8 (D) Chemical structure of loratadine (E) Effects of both compound 8 and loratadine on rabbit blood hemolysis are assessed. (F) Effects of both compound 8 and loratadine on sheep blood hemolysis are assessed. In panels A,B, E, and F, mean percent hemolysis is displayed from three biological replicates. Error bars represent the standard error of the mean. Two-way ANOVAs were used to assess variance between means of every sample. Details of this analysis are shared in Tables S1-S5.

Recently, we analyzed both compound 8 and loratadine in the absence of antibiotics for anti-virulence activity. These compounds elicited strain-specific changes to hemolysis in two hospital-acquired strains of MRSA, ATCC 43300 as well as USA100. Both are staphylococcal cassette chromosome *mec* (SCC*mec*) type II. We concluded that loratadine and compound 8 both reduced alpha and beta hemolytic activity in ATCC 43300. Compound 8 elicited effects that were smaller in magnitude but similar to loratadine. However, loratadine stimulated hemolysis in the USA100 strain. Compound 8 reduced alpha and did not change beta hemolysis in this strain. We concluded that the strain-specific differences in hemolytic modulation were likely due to signaling differences occurring downstream of Stk1 inhibition and/or additional targets were being engaged.^29^

We hypothesized that these novel anti-virulence compounds may influence hemolysis in other strains of MRSA, although strain-specific regulation would be likely. Expanding our analyses to include community-acquired strains of MRSA allows a broader view of our compounds’ effects on hemolysis across different strains and SCC*mec* types. Community-acquired strains of MRSA are known to behave differently than hospital-acquired strains, particularly with respect to regulation of antibiotic resistance and virulence genes.^34–36^ It has been postulated that community-acquired strains of MRSA are able to infect and kill heathy individuals precisely because of their higher expression of virulence factors, including hemolysins and leukocidins.^35, 36^ We chose to evaluate our compounds’ effect on hemolysin production in strains of USA300 (SCC*mec* type IV). In addition to being the most common strain of community-acquired MRSA in the United States,^37^ the Nebraska Transposon Mutant Library (NTML) was constructed using USA300 JE2.^38^ This set of valuable genetic tools allowed us to further investigate the consequences of Stk1 disruption on MRSA hemolysis. These results provide a more thorough mechanistic insight into compound 8 and loratadine’s functions as anti-virulence compounds.

## Results

### Stk1 regulation of hemolysis varies even within the same genetic lineage

We tested two different USA300 strains to begin our qualitative analysis of Stk1 modulating hemolysis. Rabbit blood agar, which reports primarily on alpha hemolysin activity, revealed that strain BAA-1556 had a larger hemolysis zone when *stk1* was deleted compared to WT (Figure 1). This indicated that Stk1 repressed alpha hemolytic activity. Note that the zone of clearing immediately surrounding the bacterial spot was easier to define than the more diffuse, outer zone. The sheep blood agar, reporting primarily on beta hemolytic activity, showed a very small degree of bruising in both WT and Δstk1 to a similar extent. Some limited alpha hemolysin activity^39^ is still observed (Figure 1). As beta hemolysin is a hot-cold hemolysin,^40, 41^ further incubation of the sheep blood agar at 4 °C revealed a larger hemolysis zone for Δstk1 compared to WT. This indicated that Stk1 may also repress beta hemolytic activity in this strain.

The second USA300 strain we tested was JE2. This strain is the genetic background for the NTML. The JE2 strain showed a different phenotypic change when *stk1* was disrupted compared to the BAA-1556 strain. Compared to WT JE2, Tn::stk1 had a similarly sized hemolysis zone on rabbit blood agar (Figure 1). JE2 strains again showed very little beta hemolytic activity via this blood agar assay. Incubation at 4 °C revealed slightly less hemolysis by Tn::stk1 compared to WT. This indicates that Stk1 may enhance beta hemolytic activity in this strain. Notably, even within the same lineage (USA300), differences in Stk1-regulated hemolysis were detected.

### Loratadine lowers hemolytic activity in both Stk1-dependent and -independent fashions through multiple Sar proteins

To more quantitatively assess the impact of Stk1 on hemolysis, liquid blood assays were performed using the WT JE2 strain as a control compared to Tn::stk1. Supernatants from Tn::stk1 cells showed a reduction in alpha hemolysin activity (94.5% the activity of Triton-X controls compared to 96.3% in WT). Using the transposon mutant of the cognate phosphatase, Tn::stp1, did not significantly change alpha hemolysin activity from WT levels (Figure 2A).

Consistent with the qualitative blood agar assays, beta hemolysin activity was very low in each JE2 strain (Figure 2B). The WT cell supernatants displayed on average 34.0% of the activity compared to Triton-X controls. Disrupting *stk1* or *stp1* did not have a statistically significant effect on beta hemolysin activity. Together, these results supported Stk1 promoting alpha but not beta hemolytic activity in USA300 strain JE2. Stp1 did not seem to influence either alpha or beta hemolytic activity.

We next determined if compound 8 and loratadine (Figure 2C-D), putative Stk1 inhibitors, would influence hemolysis in USA300 JE2. Untreated WT cultures showed 96.3% alpha hemolysin activity, while adding compound 8 (25μM) to the cultures resulted in 95.9%. Adding loratadine (50μM) decreased alpha hemolysin activity down to 78.1% (Figure 2E). We further hypothesized that if anti-virulence compounds required Stk1 to elicit these changes in hemolysis, then treating the transposon mutant with these compounds would fail to show a change in hemolysis. Strikingly, in Tn::stk1, hemolysis was further decreased by loratadine treatment. The percent hemolysis dropped to only 28.3%. This drop was not observed in Tn::stp1. A two-way analysis of variance (ANOVA) was used to test effects of these genes (WT vs transposon mutant) and loratadine (untreated vs treated) on hemolysis. The analysis revealed that there was a statistically significant interaction effect between the genetic factor (*stk1* gene disruption) and the treatment factor (loratadine), meaning the magnitude of loratadine’s effects was dependent on *stk1’s* presence (Tables S1-S3). Therefore, these results did not support loratadine influencing alpha hemolytic activity through Stk1 inhibition alone. Even when *stk1* was disrupted, alpha hemolytic activity was repressed. It remained possible that loratadine was also inhibiting one or more downstream *S. aureus* proteins, which ultimately resulted in the changes in hemolysis.

As shown in Figures 1 and 2B, the level of hemolysis using untreated samples was lower in sheep blood compared to rabbit blood. We detected consistent, significant decreases in beta hemolysin activity with loratadine treatment compared to untreated samples (Table S1, S4-S5). The magnitude of decrease (approximately 30%) was similar regardless of whether *stk1* or *stp1* were disrupted (Figure 2F) and accordingly, there were no statistically significant interaction effects of disrupting either gene with loratadine treatment (Table S1, S4-S5). More subtle decreases in beta hemolytic activity were observed when including compound 8 (Figure 2F), which is consistent with what we recently reported in other strains of MRSA.^29^ Given these limited impacts on hemolysis, results of compound 8’s effects in additional transposon mutants are presented in Figures S1-S5 and Table S6. Together, these results suggest that loratadine can inhibit both alpha and beta hemolytic activity in USA300 JE2 in a manner that is independent of Stk1 and Stp1. Additionally, we found that treating bacterial cultures with loratadine led to greater repression of hemolysis than disrupting either *stk1* or *stp1*. We previously reported Stk1 as the likely target of loratadine based on deletion mutant experiments examining loratadine-induced antibiotic potentiation.^32^ In light of that, there are at least three possibilities to explain these outcomes: 1) loratadine inhibits one or more protein(s) in addition to Stk1, 2) the disruption of the *stk1* gene by a transposon results in downstream effects unique from what occurs when Stk1 is present but its kinase activity is inhibited, or 3) disruption of the *stk1* gene results in downstream effects unique from what occurs when it is completely deleted.

To test the possibility that loratadine was impacting one or more regulators of hemolysis aside from Stk1, we utilized additional mutants in the NTML. The *sar* regulon controls the expression of many *S. aureus* virulence factors. SarA is a global transcriptional regulator, impacting gene expression of over 100 genes involved in virulence, biofilms, and metabolism^14, 16, 42^. Both SarA and SarZ are reported substrates of Stk1.^43, 44^ Phosphorylation of SarA and SarZ by Stk1 has been shown to regulate binding to the *hla* promoter.^43^ SarA has also been shown to upregulate alpha, beta, and gamma hemolysin genes directly as well as indirectly through SarA’s control of the *agr* locus.^15^ Therefore, we performed quantitative hemolysis assays comparing WT supernatants to those from Tn::sarA and Tn::sarZ. We found that loratadine no longer reduced alpha hemolytic activity of Tn::sarA bacterial supernatants (Figure 3A). This suggested that SarA might promote loratadine’s reduction in hemolysis, although interaction effects were not statistically significant (Tables S1, S7). Like the WT strain, Tn::sarZ still had reduced activity when loratadine was included (81.1%) (Figure 3A, Tables S1, S8). This indicated that SarZ is not required for loratadine’s modulation of alpha hemolytic activity. Turning to beta hemolytic activity, loratadine significantly reduced activity in WT supernatants. But when *sarA* was disrupted, both untreated and treated cells displayed background levels of beta hemolytic activity. Two-way ANOVAs revealed a very significant interaction effect between loratadine treatment and *sarA* disruption (Tables S1, S9). Disrupting *sarZ* resulted in beta hemolysin activity levels more similar to WT (Figure 3B, Tables S1, S10). This experiment supported SarA being critical in loratadine’s modulation of beta hemolysin activity.

**Figure 3:**
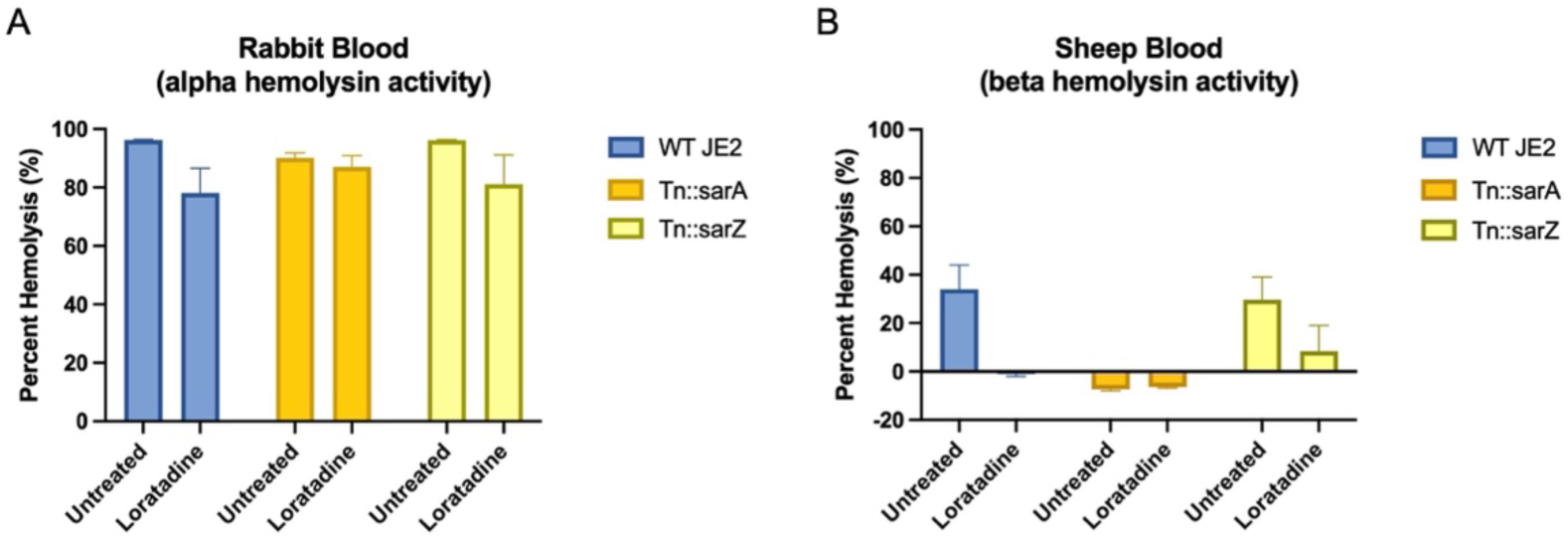
Quantitative hemolysis assays using (A) rabbit or (B) sheep blood reveal SarA impacting loratadine-induced reduction of beta hemolytic activity in USA300 strain JE2. Mean percent hemolysis is displayed from three biological replicates. Error bars represent the standard error of the mean. Two-way ANOVAs were used to assess variance between means of every sample. Details of this analysis are shared in Tables S1 and S7-10.

Repeating these assays with mutants for other SarA family genes revealed several interesting results. Disruption of *sarT* and *sarV* resulted in alpha hemolytic activity trends most similar to WT, with percent hemolysis remaining high without treatment but lower with loratadine treatment (Figure 4). This is consistent with SarT’s known role in repressing alpha hemolysin.^45^ Disrupting *sarS* resulted in the most drastic reduction in alpha hemolytic activity. Notably, SarS was originally described as a repressor of *hla* expression.^46^ This conclusion was dependent on the absence of SarA. Therefore, consistent with previous experiments by other groups,^47^ disruption of *sarS* alone may not necessarily elevate *hla* levels. Reductions in alpha hemolysin activity were also observed when disrupting *sarU* and *sarX*. These gene factors depended significantly on loratadine’s presence (Table S1, S11-14). This indicates that SarU and SarX are important in inhibiting loratadine’s repression of alpha hemolytic activity. When measuring beta hemolytic activity, all *sar* mutant supernatants showed less activity than WT, consistent with their importance in this exotoxin’s function. Statistical testing for interaction effects between gene disruptions and loratadine treatment showed that Tn::sarS and Tn::sarT had significant interaction effects (Figure 4B, Table S1, S16-20). This indicates that SarS and SarT are also important for loratadine’s repression of beta hemolytic activity. While we do not have evidence that loratadine directly interacts with these Sar proteins, the data does suggest that they impact loratadine’s effects on hemolysis directly, indirectly, or synergistically.

**Figure 4:**
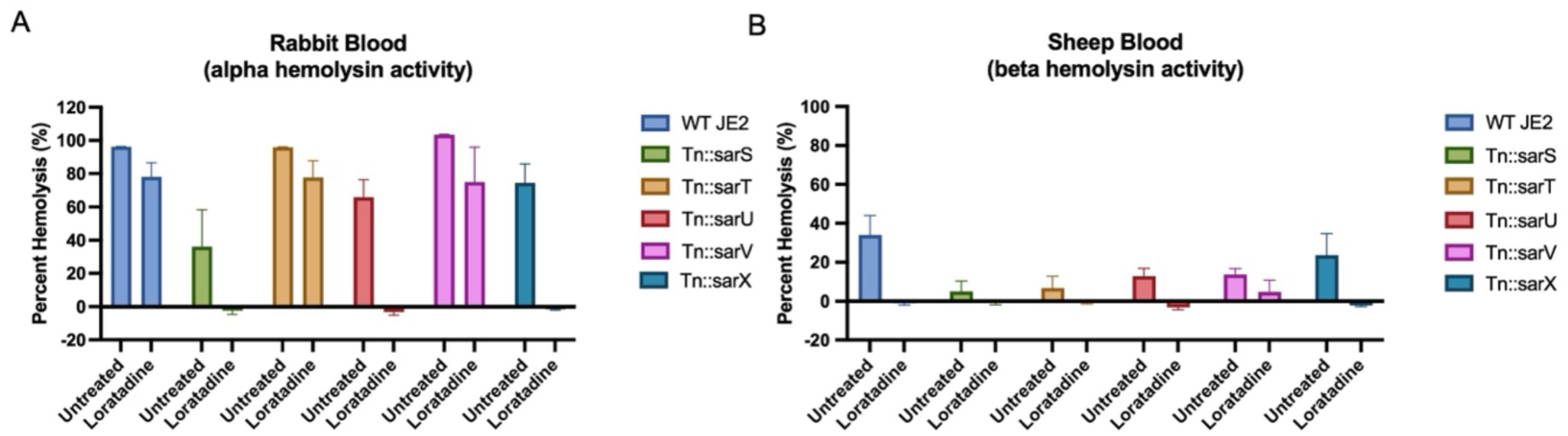
Quantitative hemolysis assays using (A) rabbit or (B) sheep blood reveal SarU and SarX inhibiting loratadine’s reduction of alpha hemolytic activity in USA300 strain JE2. SarS and SarT impact loratadine’s reduction of beta hemolytic activity. In both panels, mean percent hemolysis is displayed from three biological replicates. Error bars represent the standard error of the mean. Two-way ANOVAs were used to assess variance between means of every sample. Details of this analysis are shared in Tables S1 and S11-20.

Other known regulators of *hla* expression include the two-component system (TCS) SaeRS.^22^ These two proteins are involved in a signaling cascade that begins when SaeS, the sensor histidine kinase, detects a signal from the environment and autophosphorylates. The phosphoryl group is transferred to SaeR, the response regulator. Phosphorylated SaeR then binds to sequences in target genes, regulating their transcription. Both members of this TCS have been shown to promote *hla* expression.^48–52^

In this experiment, both Tn::saeR and Tn::saeS untreated cell supernatants showed statistically significant drops in alpha hemolysin activity compared to WT (Figure 5A, Tables S1, S21-22). Disrupting *saeR* resulted in average hemolysis levels of 27.6%, while disrupting *saeS* revealed levels that were essentially at background. This is consistent with previous groups’ reports that *saeR* or *saeS* transposon mutants have inhibited alpha hemolysin mRNA levels.^21, 23^ Once loratadine was included, hemolysis levels were at background. With alpha hemolysin activity abolished, we were unable to discern loratadine’s effects in these transposon mutants. (Table S1, S21-22). Beta hemolysin activity was not significantly reduced by *saeR* disruption, but it was by *saeS* disruption. Again, loratadine significantly lowered hemolysis activity to background levels in both transposon mutants. This also occurred in the WT strain (Figure 5B, Tables S1, S24).

**Figure 5:**
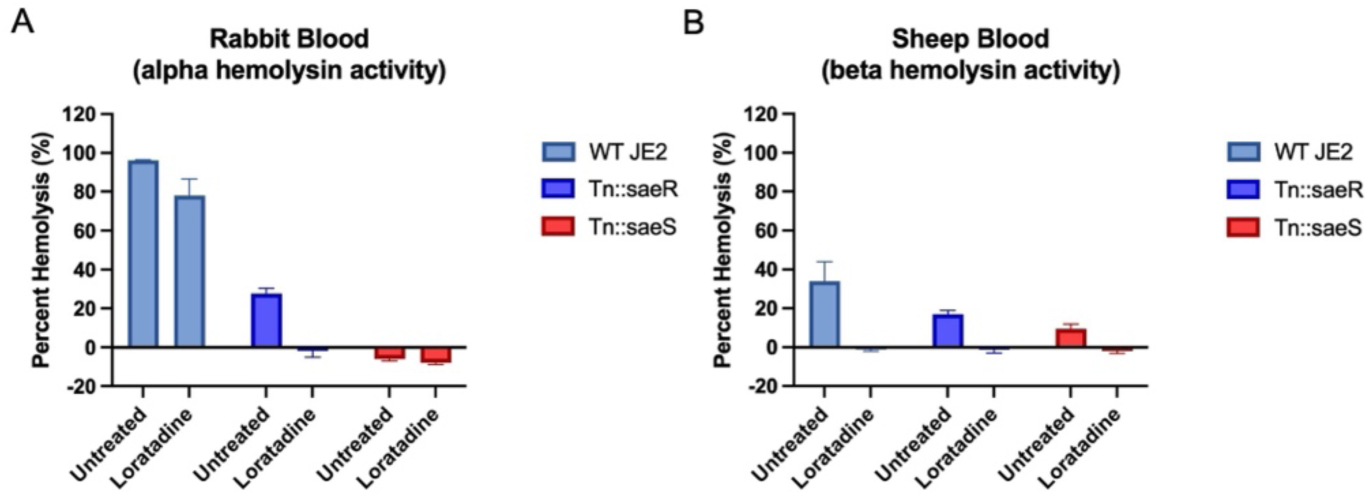
Quantitative hemolysis assays using (A) rabbit or (B) sheep blood reveal SaeRS is not required for loratadine’s reduction in alpha or beta hemolysin activity in USA300 strain JE2. In both panels, mean percent hemolysis is displayed from three biological replicates. Error bars represent the standard error of the mean. Two-way ANOVAs were used to assess variance between means of every sample. Details of this analysis are shared in Tables S1 and S21-24.

These results do not support the idea that SaeRS is required directly, indirectly, or synergistically for loratadine’s repression of hemolysis.

Another TCS with ties to hemolysis is ArlRS. The *arl* locus (autolysis related locus) consists of the histidine kinase ArlS and the response regulator ArlR.^53, 54^ There is no consensus as to how ArlRS regulates hemolysis or if it does so only in some strains of *S. aureus.*^55^ In our experiments with USA300 JE2, the Tn::arlR strain showed a very similar alpha hemolysin activity trend compared to WT, while the Tn::arlS strain revealed a larger decrease in activity with loratadine treatment compared to WT (Figure 6A). The disruption of *arlS* showed significant interaction effects with loratadine treatment, pointing to a signaling cascade where this kinase typically blocks loratadine from repressing alpha hemolytic activity (Tables S1 and S25-26). Loratadine treatment showed significant reductions in beta hemolytic activity that was not drastically different in either mutant strain compared to WT (Figure 6B, Tables S1 and S27-28). These results support ArlS being important in loratadine’s reduction in alpha hemolytic activity.

**Figure 6:**
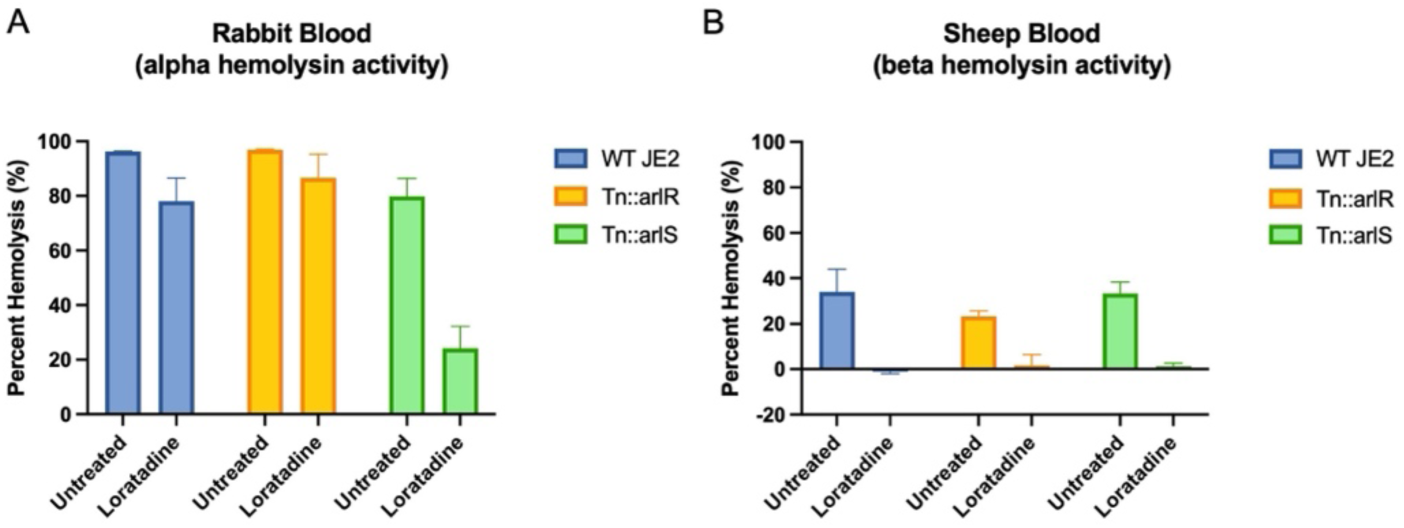
Quantitative hemolysis assays using (A) rabbit or (B) sheep blood reveal ArlS inhibiting loratadine’s reduction of alpha hemolytic activity in USA300 strain JE2. In both panels, mean percent hemolysis is displayed from three biological replicates. Error bars represent the standard error of the mean. Two-way ANOVAs were used to assess variance between means of every sample. Details of this analysis are shared in Tables S1 and S25-28.

Whether that occurs directly or indirectly through other regulatory pathways remains to be determined.

Finally, we tested the hemolysin regulator CodY. This protein has been shown to repress levels of *hla, hlb,* and *hld* in *S. aureus*.^56, 57^ Loratadine’s downregulation of alpha hemolysin activity was significantly enhanced in Tn::codY compared to WT (Figure 7A, Tables S1 and S29). Beta hemolysin activity reached similar levels compared to WT (Figure 7B, Tables S1 and S30). This suggests that CodY is another hemolysin regulator that typically inhibits loratadine’s downregulation of alpha hemolysin activity.

**Figure 7:**
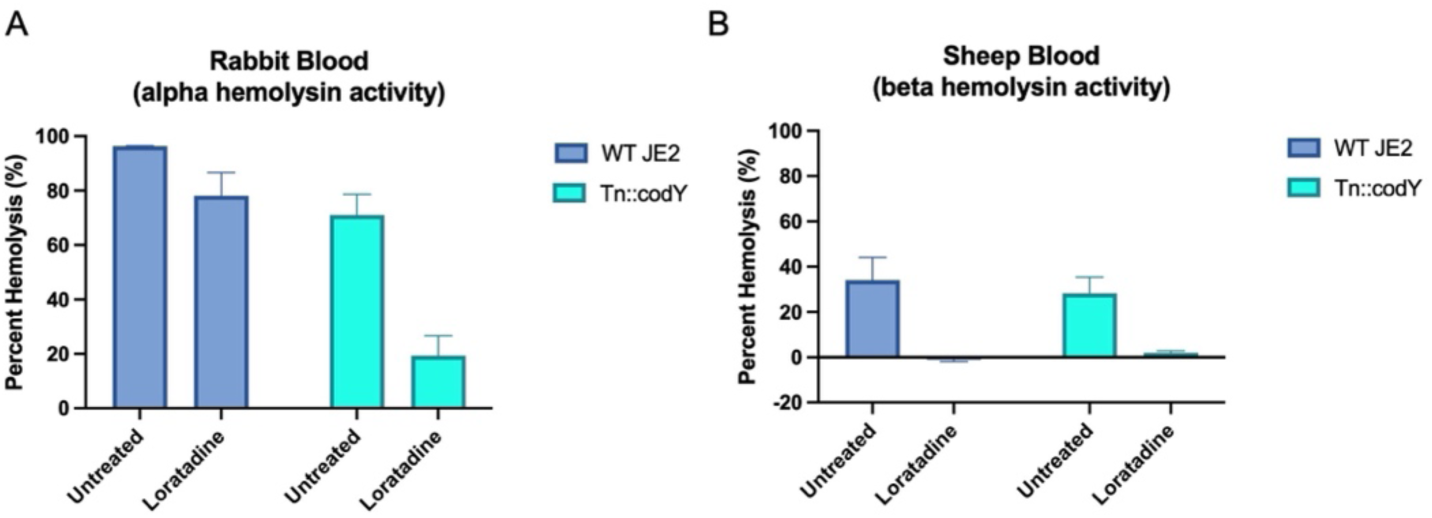
Quantitative hemolysis assays using (A) rabbit or (B) sheep blood reveal CodY inhibiting loratadine’s reduction of alpha hemolytic activity in USA300 strain JE2. In both panels, mean percent hemolysis is displayed from three biological replicates. Error bars represent the standard error of the mean. Two-way ANOVAs were used to assess variance between means of every sample. Details of this analysis are shared in Tables S1 and S29-30.

Together, these quantitative hemolysis assays using the NTML supported loratadine reducing primarily alpha hemolytic activity via a wide variety of hemolysin regulatory proteins. This novel result stems from a yet unknown mechanism but reveals that loratadine can effectively reduce hemolysis in a pleiotropic manner through both Stk1-dependent and independent actions.

### Loratadine modulates multiple cytotoxin genes at the mRNA level

To offer additional clues to loratadine’s mode of action, we utilized RNA sequencing of untreated and treated MRSA cultures. Measuring mRNA levels transcriptome-wide would help determine if loratadine was disrupting hemolysis at the transcript level or if it was limited to a post-transcriptional level. Based on our previous experiments with MRSA strains ATCC 43300 and USA100,^29^ we hypothesized that loratadine would disrupt expression of multiple hemolysins, attributing to the phenotypic effects we had observed. USA300 BAA-1556 was treated with loratadine or solvent (DMSO) for one hour before total RNA purification and sequencing. Among the hundreds of differentially expressed genes (DEGs), 377 were significantly upregulated and 324 were significantly downregulated with loratadine compared to the untreated controls (Figure 8A, Table S31). We looked for significantly enriched gene ontology (GO) categories and KEGG pathways to give biological insight to these DEGs. There was significant enrichment of DEGs in multiple translation categories (Tables S32-34) as well as the KEGG ribosomal pathway (Tables S35-37). This was consistent with loratadine’s transcriptomic impact we reported in strain ATCC 43300.^58^ While we did not find significantly enriched hemolysin-containing KEGG pathways (bacterial secretion system, transporters, or bacterial toxins), we were able to identify loratadine-induced gene expression changes in every hemolysin gene *S. aureus* expresses. Figure 8B-G shows that loratadine treatment resulted in downregulation of *hla* and *hld*, while *hlb*, *hlgA*, *hlgB*, and *hlgC* were upregulated under these conditions. While only *hld* downregulation was statistically significant, this in vitro experiment used cultures that were treated with loratadine for one hour, in contrast to 24 hours needed to optimally detect excretion of these toxins in quantitative hemolysis assays. Since gamma and delta hemolysins are also considered leukocidins, we examined the RNA-seq dataset for additional toxins that loratadine could be modulating. As shown in Figure 8H-L, all of the PSMs detected in this strain of MRSA were significantly downregulated by loratadine. Several of these hemolysin gene expression changes were validated in bacterial cultures treated with loratadine that were independent from those used in RNA-seq. The results determined by RT-qPCR confirmed that *hla* is downregulated, and *hlgA* and *hlgC* are upregulated by loratadine (Figure S6). Together, these results supported loratadine disrupting a plethora of cytotoxin gene expression events in hypervirulent USA300.

**Figure 8:**
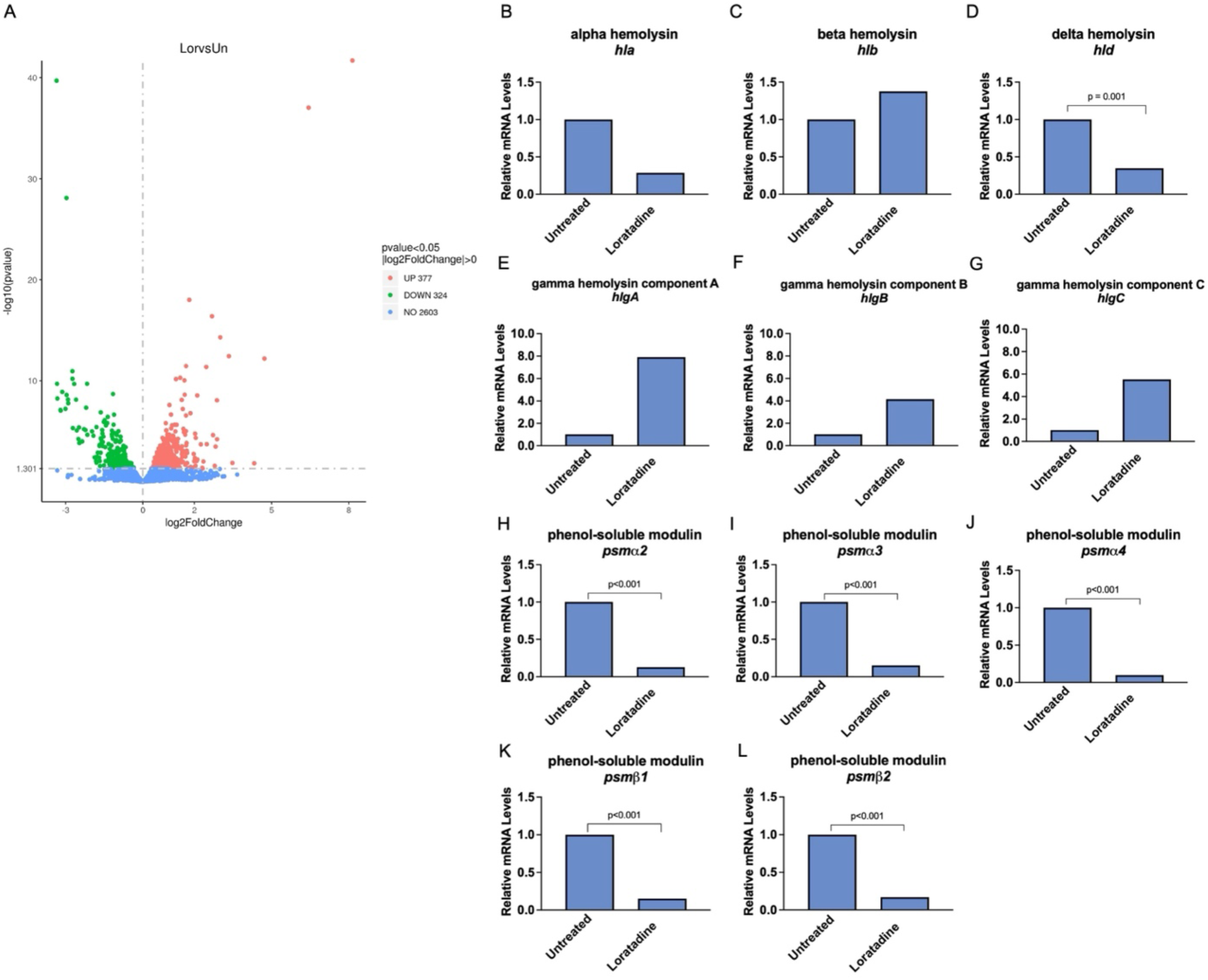
RNA-seq reveals that loratadine modulates multiple cytotoxins at the mRNA level. (A) A volcano plot showing the magnitude of downregulated genes (green) and upregulated genes (red) in loratadine compared to untreated samples. Results show average mRNA levels of *hla* (B), *hlb* (C), *hld* (D), *hlgA* (E), *hlgB* (F), *hlgC* (G), *psmα2* (H), *psmα3* (I), *psmα4* (J), *psmβ1* (K), and *psmβ2* (L) from biological triplicate samples after loratadine treatment relative to the untreated control samples. Adjusted p values ≤0.05 are displayed.

### Loratadine impacts expression of global virulence regulators *agrBDCA* and *vraSR*

Given that loratadine, a likely Stk1 inhibitor, was affecting expression of multiple cytotoxin genes, we next looked at the complex regulatory network controlling these virulence factors. We hypothesized that one or more top-level effectors could be modulated at the mRNA level with loratadine treatment. The consequence of this modulation would explain the widespread disruption in downstream cytotoxin expression we detected.

The *agr* locus is a major regulator of virulence and is considered the main regulator of alpha hemolysin expression. It facilitates this regulation via RNAIII as an effector RNA molecule which upregulates expression of both *hla* and *hlb.* ^5, 13, 14^ RNAIII also prevents translation of repressor of toxin (Rot), leading to upregulation of multiple toxins. ^59–61^ The *agr* regulon also directly controls PSMs in a positive manner.^62^ Furthermore, as we reported here that *hld* (transcribed within RNAIII) was significantly downregulated by loratadine, we examined whether any differential gene expression was occurring to *agr* genes themselves (*agrBDCA*). As shown in Figure 9A-D, the expression of each gene in this locus was lowered by loratadine to approximately half the levels of untreated samples (Figure 9, Table S31). Next, we looked to the SarA protein family genes including MgrA and the Sar genes examined in Figures 3 and 4. None of these hemolysis regulators had their mRNA levels significantly impacted by loratadine (Table S31). Additional hemolysin regulatory genes such as *codY*, *saeRS*, and *arlRS* also showed no significant changes to gene expression. The mRNA levels of *stk1* and *stp1* were also unaltered compared to untreated samples (Table S31). Together, this high throughput analysis revealed that loratadine treatment resulted in lowered mRNA levels of the entire *agr* system, likely contributing to the reduction in hemolysin and PSM mRNA levels we detected.

**Figure 9:**
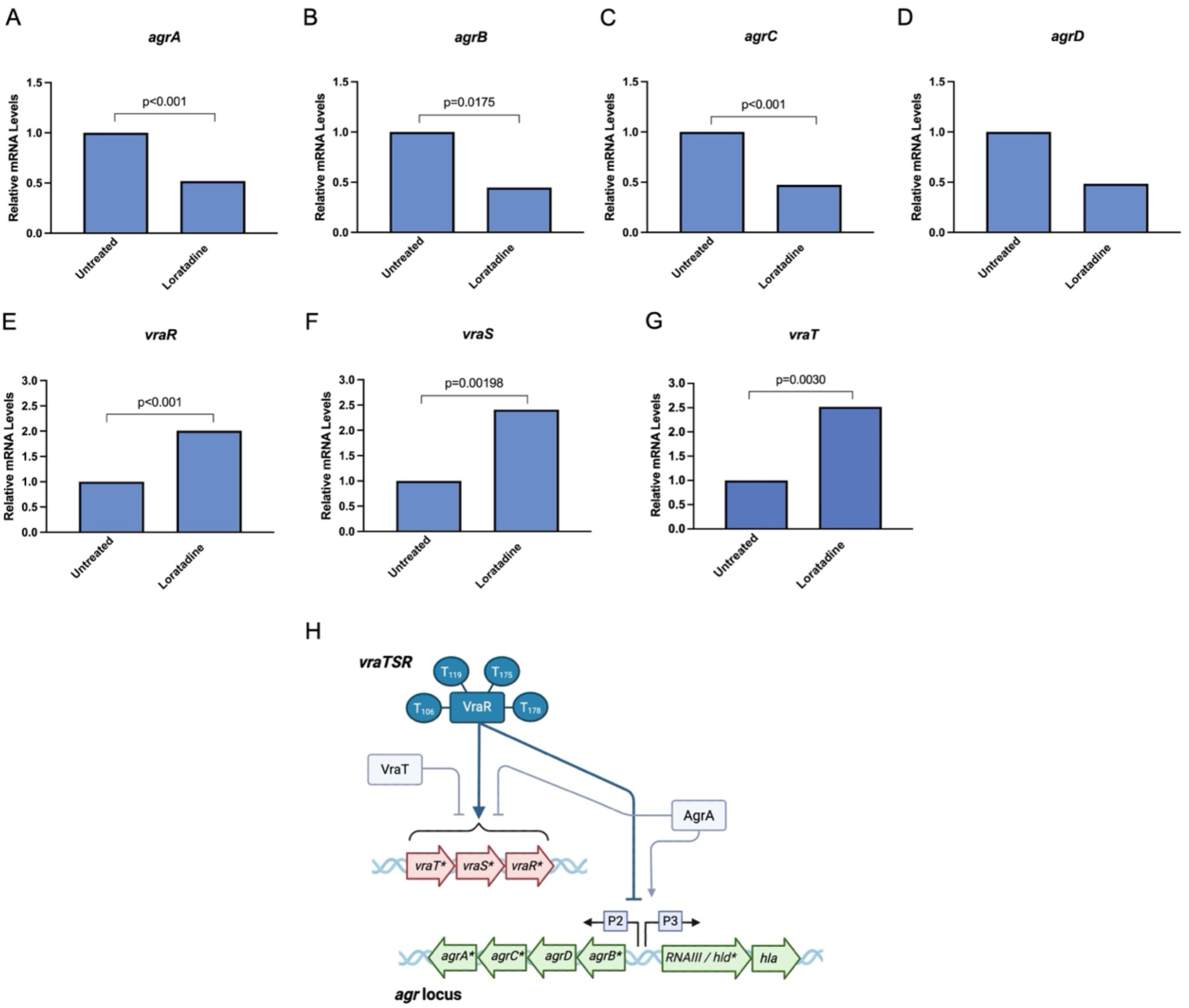
RNA-seq reveals that loratadine downregulates mRNA levels of the *agr* locus and upregulates *vraTSR*. Results show average mRNA levels of *agrA* (A), *agrB* (B), *agrC* (C), *agrD* (D), *vraR* (E), *vraS* (F), and *vraT* (G) from biological triplicate samples after loratadine treatment relative to the untreated control samples. Adjusted p values ≤0.05 are displayed. (H) Graphic of loratadine’s impacts to these two regulatory loci. Genes are noted as arrows denoting whether the gene is found on the plus strand (arrow to the right) or minus strand (arrow to the left). Genes colored gray had no difference in expression between loratadine treated and untreated samples via RNA-seq. Genes colored green were downregulated with loratadine treatment as compared to the untreated control. Genes colored red were upregulated with loratadine treatment as compared to the untreated control. An asterisk indicates an adjusted p value of < 0.05. Key proteins involved in cytotoxin regulation have been shown as rectangles. Proteins colored in teal are substrates of Stk1 and the known phosphorylation sites are indicated as circles with the 1-letter amino acid abbreviation. Regulatory pathways that depend on phosphorylation by Stk1 are shown in teal lines. Pathways that do not directly depend on phosphorylation by Stk1 are shown in light blue lines.

*S. aureus* must not only sense host heme but respond to its levels by utilizing the heme-sensing TCS HssSR. The sensor component is a histidine kinase encoded by *hssS*, while the regulatory protein is encoded by *hssR*. This TCS coordinates with a heme regulated transporter efflux pump HrtAB to avoid intracellular heme toxicity.^63^ To investigate loratadine potentially modulating these heme homeostasis players, we examined RNA-seq data for significant gene expression changes. We did not detect loratadine modulating *hssS*, *hssR*, *hrtA*, or *hrtB* levels (Table S31). Loratadine also did not affect genes that compose the heme transport system *hts* (Table S31). Loratadine did upregulate one gene involved in the heme acquisition system, iron-regulated surface determinant *isdB* ^64, 65^ (Table S31). Finally, the three-component system *vraTSR* indirectly regulates hemolysin transcription by controlling *agr*.^66^ VraR is the response regulator and VraS is the histidine kinase originally identified for their roles in vancomycin resistance and other cell wall damaging agents.^67^ VraT (formerly known as YvqF) has a less-defined role, but is proposed to help the VraR regulon ensure optimal methicillin resistance.^66^ VraR has been shown to bind the promoter of *agr* and inhibit the function of this system.^68^ VraR has also been identified as an Stk1 substrate, where phosphorylation of VraR reduces or abolishes its DNA binding activity to its own promoter.^69^ Likewise, AgrA has been recently shown to bind and inhibit the *vraTSR* promoter, further entwining their relationship in hemolysis regulation.^70^ In our RNA-seq experiment, loratadine resulted in upregulation of this three-component system to approximately twice the levels of untreated samples (Figure 9E-G, Table S31).

Given loratadine’s modulation of these global hemolysis regulators at the mRNA level (Figure 9H), we next tested if loratadine’s mode of action required the VraR regulon to reduce *agr* levels. We used RT-qPCR to measure *agrA* levels in the USA300 strain JE2 as well as transposon mutants for *vraR* and *vraS*. As shown in Figure 10A, USA300 JE2 cells showed *agrA* mRNA levels reduced to approximately 0.8X the levels of untreated cells, similar to the USA300 BAA-1556 strain (Figure 9A). However, when Tn::vraR was treated with loratadine, this repression in *agrA* was no longer observed. All other tested transposon mutants (including those for *stk1* and *stp1*) showed loratadine reducing *agrA* like the WT strain (Figure 10A). We extended this mutant analysis to other differentially expressed hemolysis genes and their regulators. Consistent with the USA300 BAA-1556 strain (Figure 9A), loratadine also upregulated *vraR* levels by approximately 2X in USA300 JE2 (Figure 10B). In contrast, loratadine no longer upregulated *vraR* in the Tn::vraR mutant (Figure 10B). All other mutants in the experiment responded like the WT strain. The same trend was observed for *vraS* mRNA (Figure 10C), indicating that VraR is critical for loratadine’s mode of action. Hemolysins *hla* and *hld* were still downregulated, similarly to WT JE2, regardless of the transposon mutant used (Figure 10D,E). Together these results reveal that VraR, a known substrate of Stk1 and regulator of the Agr system, is important for loratadine’s modulation of virulence gene expression while Stk1 and Stp1 are dispensable.

**Figure 10:**
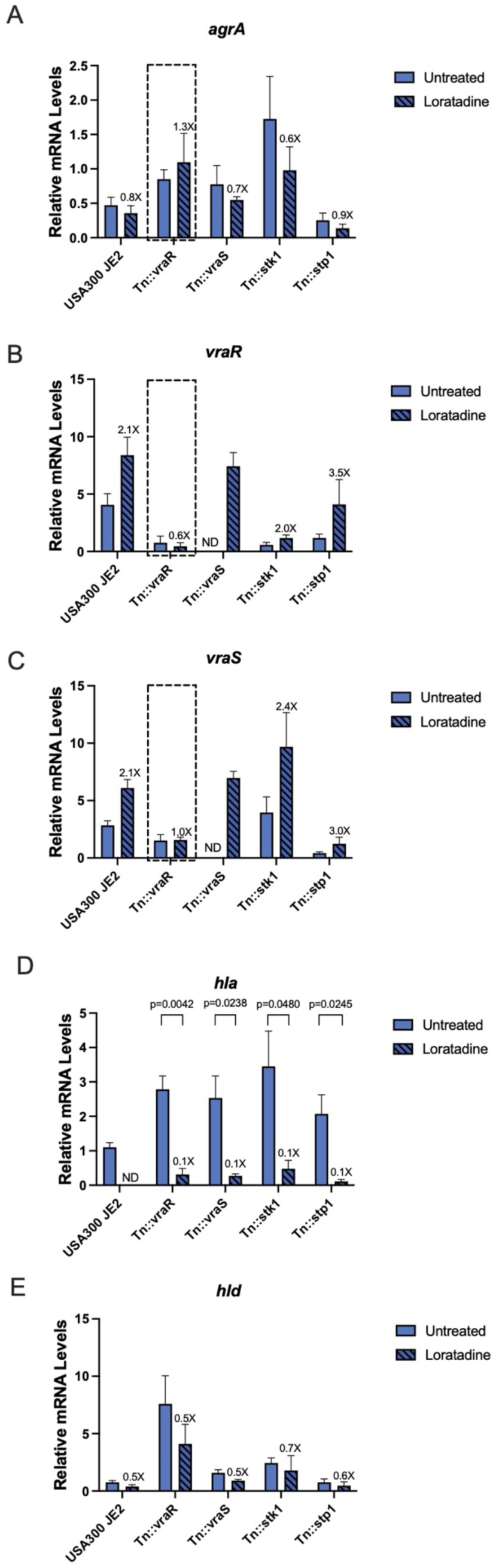
NTML analysis reveals that loratadine depends on VraR for its mode of action. WT USA300 JE2 and a series of transposon mutants were used to measure mRNA levels of *agrA* (A), *vraR* (B), *vraS* (C), *hla* (D), *and hld* (E) from biological triplicate samples after loratadine treatment relative to the untreated control samples. In all panels, mRNA levels are relative to a 16S reference gene. Error bars represent the standard error of the mean. Labels over loratadine samples indicate fold change from untreated controls. Unpaired student’s t-tests were used to assess statistically significant differences. Resulting p values ≤0.05 are displayed. ND, not detected.

Loratadine’s major impact on hemolysis through two top-level regulatory systems, *agrDCAB* downregulation and *vraTSR* upregulation, depend at least partly on VraR. It is also possible that in addition to these transcript-level changes, the hemolysis-lowering effects of loratadine are occurring at the post-translational level where the lack of VraR, SarA, and SarZ phosphorylation by the kinase Stk1 contributes to cytotoxin modulation. Regardless, loratadine’s ability to disrupt hemolysis through both Stk-1-dependent and Stk-1-independent pathways (Figure 11) makes it particularly effective in these hypervirulent strains of MRSA that utilize redundant pathways to produce cytotoxins.

**Figure 11:**
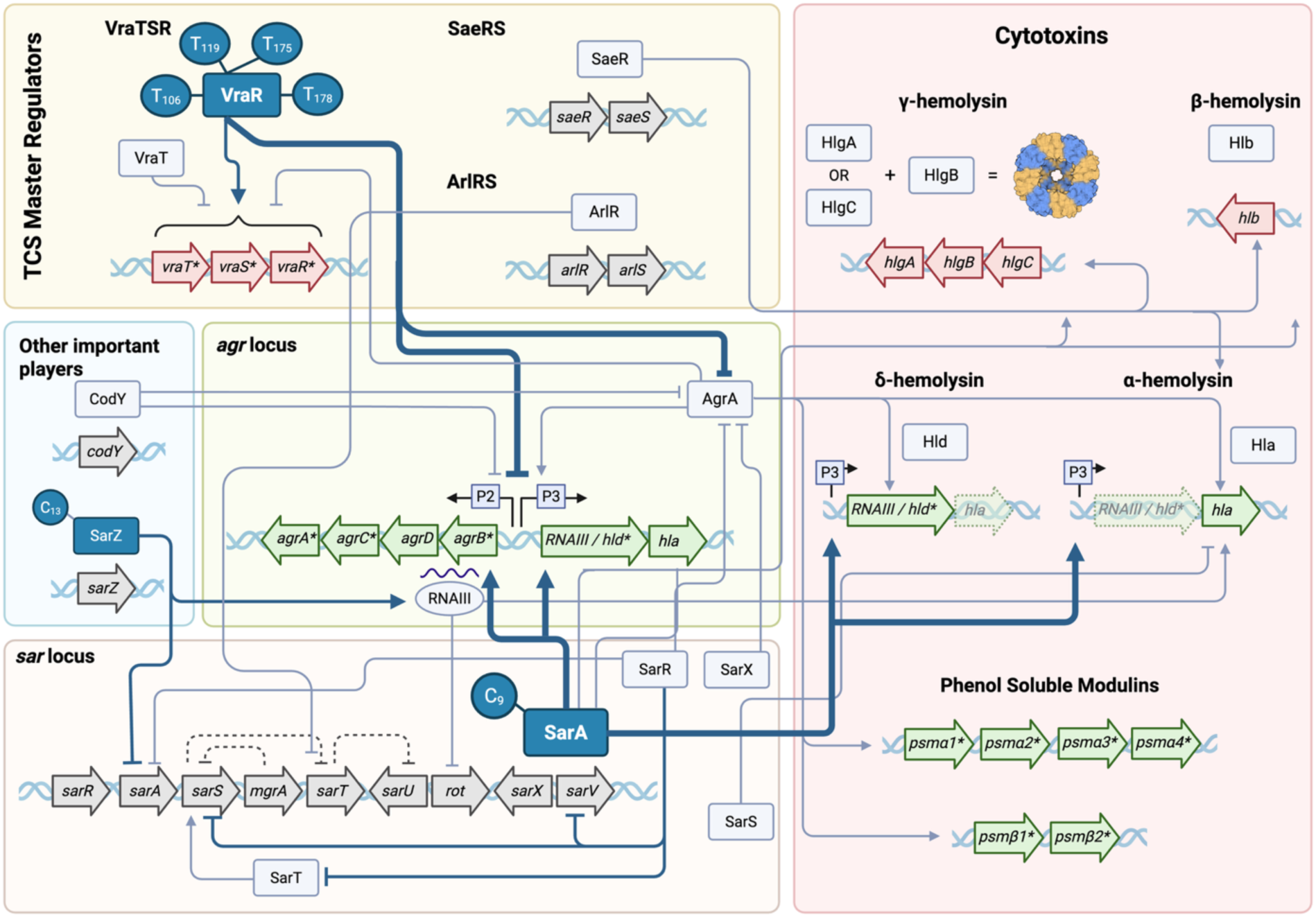
Loratadine’s influence on regulatory pathways involved in cytotoxin production in *S. aureus*. Genes are noted as arrows denoting whether the gene is found on the plus strand (arrow to the right) or minus strand (arrow to the left). Genes colored gray had no difference in expression between loratadine treated and untreated samples via RNA-seq. Genes colored green were downregulated with loratadine treatment as compared to the untreated control. Genes colored red were upregulated with loratadine treatment as compared to the untreated control. An asterisk indicates an adjusted p value of < 0.05. Key proteins involved in cytotoxin regulation have been shown as rectangles. Proteins colored in teal are substrates of Stk1 and the known phosphorylation sites are indicated as circles with the 1-letter amino acid abbreviation. Regulatory pathways that depend on phosphorylation by Stk1 are shown in teal lines. Pathways shown in this paper to be important to loratadine’s mode of action are shown in heavily weighted teal lines. Pathways that do not directly depend on phosphorylation by Stk1 are shown in light blue lines.

### Loratadine downregulates multiple hemolysins in a human cell line infection model

While this transcriptomic experiment revealed modulation of important cytotoxins like hemolysins and their upstream regulators at the mRNA level, we next wanted to investigate loratadine in a more physiologically relevant experiment where human host cells are involved (Figure 12A). In culturing the human endothelial-like EA.hy926 cell line, we hypothesized that loratadine would not be cytotoxic, given its safety profile as an FDA-approved antihistamine. Furthermore, we have previously tested a range of loratadine concentrations in the human embryonic kidney HEK 293 cell line and showed no significant effect on cell viability.^58^ In the EA.hy926 cell line, loratadine was not more cytotoxic than the untreated (DMSO) control after 24 hours (Figure 12B). This held true, even at double the concentration (100 μM) used in RNA-seq experiments. Potential cytotoxicity was further examined in this cell line in the presence of USA300 JE2. As expected, cytotoxicity was elevated due to bacterial infection, but no significant differences were detected between any concentration of loratadine and the untreated control (Figure 12C). We did observe rounding of these endothelial-like host cells only with loratadine treatment (Figure S7A). The effect was temporary, with roundness depending on time but independent of bacterial infection (Figure S7B, C, Tables S38-39). Furthermore, it was specific to the endothelial-like cell type. Examining the morphology of epithelial human cervical carcinoma cells (HeLas) did not show enhanced roundness triggered by loratadine (Figure S7D, Table S40). This reversible cell rounding in endothelial cells has been reported previously to be attributed to a host response to histamines.^71^ Therefore, it is likely that the antihistamine, loratadine, was inducing this morphological change. Regardless, any temporary rounding of the host cells did not equate to cytotoxicity. Compound 8 was also deemed non-cytotoxic in this cell line (Figure S8A, B).

**Figure 12:**
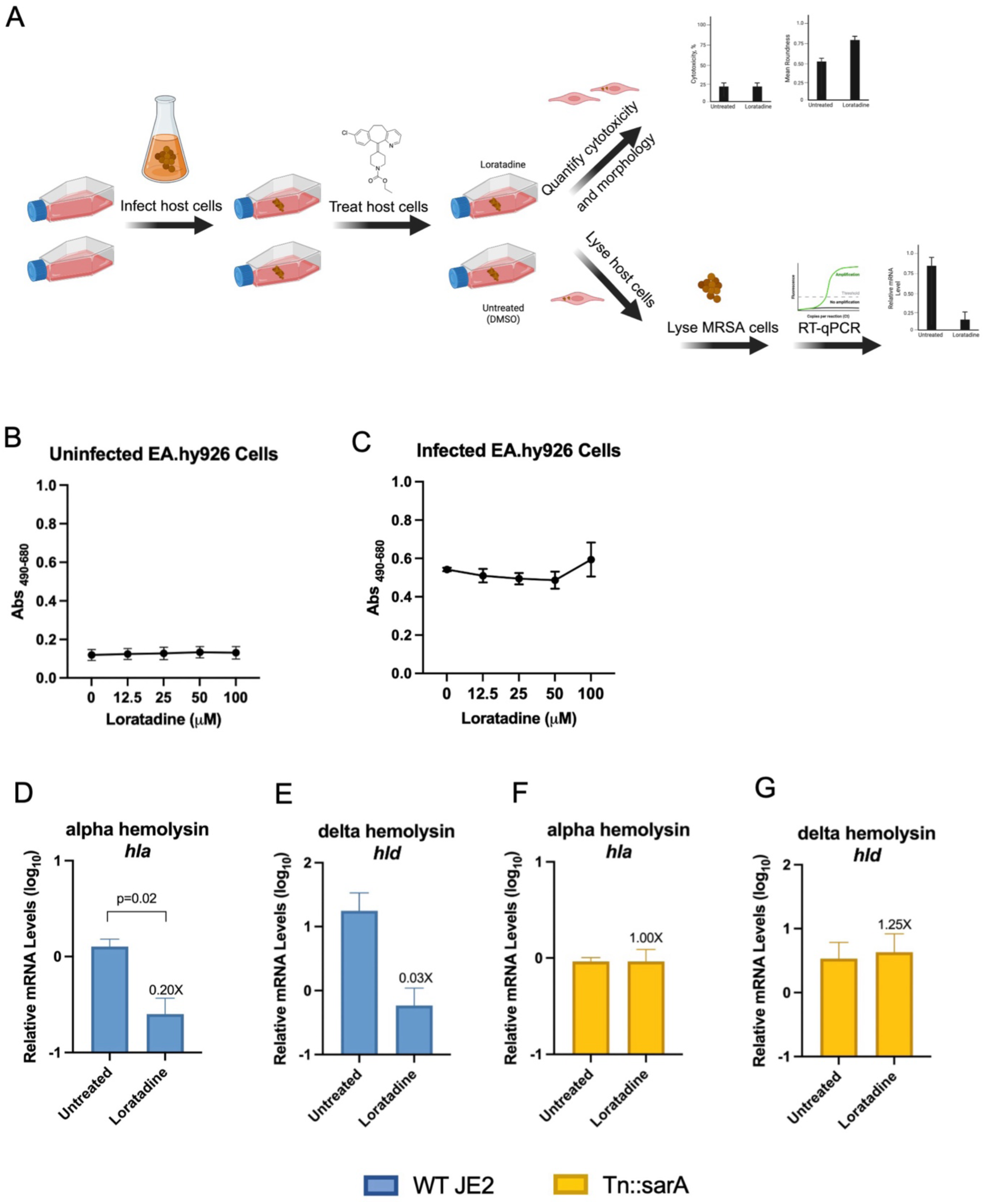
Human cell line infection model shows loratadine does not induce cytotoxicity and multiple hemolysins’ mRNA levels get suppressed. (A) Overview of human cell line infection model (B) Cytotoxicity results of loratadine in EA.hy926 cells after 24 hrs. (C) The same experiment as shown in (B), but in the presence of USA300 JE2 MRSA. Error bars represent the standard error of the mean. RT-qPCR results of (D) *hla* and (E) *RNAIII*/*hld* levels in internalized USA300 JE2 after 24 hrs. of loratadine treatment. RT-qPCR results of (F*) hla* and (G) *RNAIII*/*hld* in internalized USA300 JE2 Tn::sarA after 24 hrs. of loratadine treatment. In panels D-G, mRNA levels are relative to a 16S reference gene. Error bars represent the standard error of the mean. Labels over loratadine samples indicate fold change from untreated controls. Unpaired student’s t-tests were used to assess statistically significant differences. Resulting p values ≤0.05 are displayed.

Once loratadine was shown to be non-cytotoxic in this cell line, we used the EA.hy926 cells as a host to model a MRSA infection in humans, as described by others.^72^ We infected the host cells with USA300 JE2 then treated with loratadine at 50 μM final concentration or kept DMSO treated control flasks incubating for 24 hours. MRSA that had invaded the human host cells were isolated, RNA was purified, and RT-qPCR was used to measure mRNA levels within the internalized bacteria (Figure 12A). As shown in Figures 12D and E, loratadine dramatically reduced the levels of both *hla* and *RNAIII/hld*. These reductions in multiple key virulence factors were even more pronounced than in in vitro experiments (Figure 8). Additional virulence factors *hlgA* and *hlgC* were not consistently detected above background in these infection model conditions (Table S41). This indicates that at least some of loratadine’s anti-virulence effects stem from changes at the gene expression level that still occur in a human cell line infection model.

When USA300 JE2 Tn::stk1 or Tn::stp1 was used in the infection model, no hemolysin genes were amplified, while a reference gene was (Table S41, Figure S9). The same experimental procedures and reagents were used as in the WT USA300 JE2 experiments, which resulted in amplification of both *hla* and *RNAIII*/*hld.* Therefore, it was most likely the disruption of *stk1* and *stp1* that caused this lack of expression under these conditions. This was consistent with work published in other reports ^29, 30^ along with data shown in Figures 1 and 2 where Stk1 promotes expression of multiple hemolysin genes. Since Stp1 dephosphorylates Stk1,^24^ it is reasonable that disruption of either regulatory protein heavily disrupts hemolysin gene expression. This lack of expression of multiple hemolysin genes in these transposon mutant strains prevented us from determining if either regulatory protein was required for loratadine’s effects on hemolysis mRNA levels in the infection model.

Finally, we were able to examine a key Stk1 substrate that has established, global roles in regulating multiple hemolysins: SarA. Not only is SarA an Stk1 substrate, but its disruption resulted in the most statistically significant impact to loratadine’s reduction in beta hemolysin activity reported here (Figure 3B, Table S1) while also resulting in alpha hemolysin activity modulation (Figure 3A). Additionally, examining SarA’s role allowed us to more specifically investigate hemolysin expression in this virulence model as compared to other loratadine-affected regulators such as VraR. Our concern was that *vraR* disruption would result in impact to multiple pathways related to infection, antibiotic resistance, and stress. Using Tn::sarA in the infection model, we found that loratadine’s reduction of both *hla* and *RNAIII*/*hld* levels was abolished (Figure 11F, G). These *hla* results are consistent with the alpha hemolysin activity abrogation in the Tn::sarA hemolysis assay (Figure 3A). These results support a model where loratadine relies on SarA for triggering this reduction in key virulence factors during a MRSA infection.

## Discussion

Community-acquired MRSA (CA-MRSA) strains are generally considered to be hypervirulent due to their ability to infect otherwise healthy individuals without any obvious connections to health-care centers. It is believed that these hypervirulent strains employ numerous, redundant, interacting regulatory systems to control virulence, infection, and survival in hosts and the environment. We have previously reported that loratadine has a pleiotropic effect in MRSA at the mRNA level, suggesting that it likely inhibits multiple regulatory proteins either directly or indirectly through disruption of regulatory pathways.^58^ Based on these prior observations, we sought to determine whether loratadine could exert control over a common feature of hypervirulent, CA-MRSA strains: hemolysis. USA300 is the most common strain of CA-MRSA in the United States and produces high levels of alpha hemolysin via elevated expression of *agr*.^73^ We report the novel result that loratadine treatment can reduce the mRNA levels of *agr* and this is partly dependent on VraR, a known Stk1 substrate. This mode of action likely includes downstream impacts on other virulence genes and processes in addition to cytolysis. We also show that loratadine’s hemolysis modulation involves multiple proteins in the SarA family (themselves global hemolysin regulators). Therefore, it is not surprising that a drug that displays both Stk1-dependent and independent effects widely impacts hemolysis. In addition to the global virulence regulators tested in this study, MgrA (a SarA family member) has also been implicated as a target of loratadine.^33^ MgrA is a known regulator of *hla,* both directly and through additional regulators including *spa* and the *agr* locus. ^74, 75^ Unfortunately, the Tn::mgrA mutant is not available as part of the NTML and was therefore unable to be similarly assessed in these studies. Finally, we show that loratadine’s downregulation of hemolysin gene expression is enhanced in a human cell culture infection model with substantial reductions in both *hla* and *RNAIII*/*hld* levels. These reductions depended on the presence of SarA. These results show interference with production of multiple hemolysins and further support loratadine being an efficacious anti-virulence drug.

Loratadine is already FDA-approved and safe for use in both children and adults. Furthermore, it has been shown to be safe at levels above what is commonly prescribed for allergic rhinitis.

Adding to the benefits of loratadine is its displayed polypharmacy. Both our group and others have shown that loratadine likely inhibits (directly or indirectly) multiple *S. aureus* regulatory proteins.^33, 58^ This attribute affords loratadine the possibility of reducing virulence in a pleiotropic manner that *S. aureus* will not be able to easily avoid via mutation.

Studying hemolysis by bacterial pathogens has its challenges. Not only does *S. aureus* encode a collection of hemolysin genes, but the phenotypic effects of these gene products often vary depending on the strain.^29^ Transcript-level effects are also strain-dependent. Loratadine’s modulation of the *agr* locus reported here did not extend to hospital-acquired USA100 or ATCC 43300, for example, yet the drug downregulated *hla* and *hld* mRNAs in those strains through other means.^29^ These strain-specific differences are important to examine before drawing conclusions about the microorganism. Temporal changes in gene expression are an additional compounding factor to consider. For example, treating bacterial cultures for one hour with loratadine offered a “snapshot” of gene expression for RNA-seq experiments, where most hemolysins were downregulated and the gamma hemolysins were upregulated (Figure 8). While informative, we recognize that mRNA levels fluctuate constantly and any experiment with a single time point is not indicative of mRNA levels at all time points.^29^ Additionally, variability between replicate hemolysis experiments can occur.^76^ We consistently encountered this primarily with sheep blood. Variability was minimized in quantitative hemolysis assays by using the same batch of rabbit or sheep blood and running biological triplicate experiments on the same 96-well plate. Importantly, by using blood sourced from two different animals, we were able to more thoroughly examine anti-virulence effects on both alpha and beta hemolysins. Our results are consistent with a previous publication using BAA-1556 treated with loratadine. This group also showed that loratadine reduced alpha hemolysin activity.^33^ This report did not test any blood other than rabbit, limiting conclusions that could be drawn to just those on alpha hemolysin activity.

This is the first report of using loratadine in a human cell line infection model. This approach takes into consideration the host factors that respond to bacterial infection. To our knowledge, very few publications include gene expression measurements of internalized *S. aureus* in an infection model.^77–79^ Most reports that examine intracellular *S. aureus* focus on enumerating or visualizing bacteria, rather than drawing important gene expression conclusions in a physiologically relevant infection model. One group using *S. aureus* in a human lung epithelial model noted the likely dramatic transcriptomic changes triggered by internalization into a host.^78^ Importantly, our results show that hemolysin suppression at the mRNA level also occurs in this infection model. Additional studies are warranted for examining loratadine repurposed from an antihistamine to an anti-virulence drug.

## Materials and Methods

### Bacterial strains

*S. aureus* 43300, USA100 (BAA-1753), and USA300 (BAA-1556) were purchased from the American Type Culture Collection (ATCC). USA300Δstk1 (ALC6630) was from Dr. Ambrose Cheung.^26^ The following reagents were provided by the Network on Antimicrobial Resistance in *Staphylococcus aureus* (NARSA) for distribution through BEI Resources, NIAID, NIH: *Staphylococcus aureus* subsp. *aureus*, Strain JE2, Transposon Mutant strain JE2, NR-46543; Tn::stk1, transposon mutant NE217 (SAUSA300_1113), NR-460; Tn::stp1, Transposon Mutant NE1919 (SAUSA300_1112), NR-48461; Tn::sarA, Transposon Mutant NE1193 (SAUSA300_0605), NR-47736; Tn::sarZ, Transposon Mutant NE567 (SAUSA300_2331), NR-47110; Tn::sarS, Transposon Mutant NE165 (SAUSA300_0114), NR-46708; Tn::sarT, Transposon Mutant NE514 (SAUSA300_2437), NR-47057; Tn::sarU, Transposon Mutant NE96 (SAUSA300_2438), NR-46639; Tn::sarV, Transposon Mutant NE1941 (SAUSA300_2218), NR-48483; Tn::sarX, Transposon Mutant NE296 (SAUSA300_0654), NR-46839; Tn::saeR, Transposon Mutant NE1622 (SAUSA300_0691), NR-48164; Tn::saeS, Transposon Mutant NE1296 (SAUSA300_0690), NR-47839; Tn::arlR, Transposon Mutant NE1684 (SAUSA300_1308), NR-48226; Tn::arlS, Transposon Mutant NE1183 (SAUSA300_1307), NR-47726; Tn::codY, Transposon Mutant NE1555 (SAUSA300_1148), NR-48097; Tn::vraR, Transposon Mutant NE554 (SAUSA300_1865), NR-47097; Tn::vraS, Transposon Mutant NE823 (SAUSA300_1866), NR-47366.

### Blood agar assays

Rabbit and sheep blood agar were purchased from VWR. Assays were performed as previously described.^29^

### Quantitative hemolysis assays

Experiments were performed as previously described ^29^ with the following changes. Incubation occurred at 37 °C in the plate reader for one hour, measuring in 5-minute intervals. The timepoint and supernatant dilution that resulted in hemolysis in the linear range of the assay were used for all replicates with that blood type. Experiments included three technical replicates and three biological replicates. Mean percent hemolysis values of biological replicates were analyzed between WT and deletion mutants using GraphPad Prism v10.3.1. One-way ANOVAs were used for comparing means of untreated samples of WT and mutant strains. These were between measurements with no repeated measures. Adjusted p-values were generated through multiple hypothesis testing using Tukey’s test. Two-way ANOVAs were used for comparing means of untreated, compound-treated, WT, and mutant strains. These were between measurements with no repeated measures. Loratadine or compound 8 treatment served as the row factor and gene deletion served as the column factor. Interaction effects were included. Adjusted p-values were generated through multiple hypothesis testing using Tukey’s test.

### RNA-seq

Bacterial cultures of USA300 BAA-1556 were grown as biological triplicates in cation-adjusted Mueller-Hinton broth (CAMHB) with shaking incubation at 37 °C overnight. Mid log-phase cultures were diluted to approximately 5 x 10^5^ CFU per mL in CAMHB. After aliquoting cultures into sterile culture tubes, loratadine was added to a final concentration of 50 μM. Cultures treated with solvent DMSO served as negative controls. Cultures were then incubated at 37 °C with shaking for 1 hour, pelleted, and frozen in a -80 °C freezer until total RNA purification. Total RNA was purified using Qiagen RNeasy kits and Invitrogen’s Turbo DNase. Quantification and purity assessment were achieved using a DeNovix microvolume spectrophotometer. Additional RNA quality control was performed by Novogene, Inc. using a BioAnalyzer (Table S42). All library construction, sequencing, processing, and bioinformatic steps were performed by Novogene, Inc. as previously described.^80^ Read quality and reference genome mapping results are shown in Tables S43 and S44. Raw sequencing and processed data were deposited to Gene Expression Omnibus (GEO) under accession number GSE291913.

### Human cell culture

EA.hy926 endothelial-like cells were purchased from the University of North Carolina at Chapel Hill Tissue Culture Facility. HeLa cells (epithelial) were purchased from ATCC. Both cell lines were cultured in Dulbecco’s Modified Eagle’s Medium (DMEM) containing 10% fetal bovine serum at 37°C and 5% CO2. Cultures routinely tested negative for mycoplasma contamination and were passaged less than 12 times.

### Cytotoxicity assays

EA.hy926 cells were assayed for cytotoxicity with varying amounts of loratadine or compound 8. Cultures treated with solvent DMSO served as negative controls. The CyQUANT LDH cytotoxicity assay kit from Invitrogen was used. The optimum cell number for accurate lactate dehydrogenase (LDH) detection was experimentally determined as detailed in the manufacturer’s instructions at 5,000 cells per well of a 96-well plate. Cells adhered with overnight incubation, followed by 24 hrs. of drug treatment. Technical triplicates were included. Results are from biological triplicate experiments. Average LDH release was calculated according to manufacturer’s instructions among biological triplicates by background subtracting the absorbance at 680nm from the absorbance at 490nm. GraphPad Prism v10.3.1 was used to generate figures and analyze statistically significant differences with a one-way ANOVA. When USA300 JE2 was included in the cytotoxicity assays, 20,000 cells per well were plated to account for cell death during infection. Infections occurred following previously published methods^72^ with the following changes. After 90 minutes of infection, stop media containing gentamycin was added for one hour to kill external MRSA cells, then cells were treated with either loratadine or compound 8-containing DMEM at the indicated concentrations, or an untreated (DMSO) control.

### Human cell line infection model

These experiments were based on previously published methods^72^ with the following changes. EA.hy926 cells were infected with MRSA cultures for 90 minutes, stop media containing gentamycin was added for one hour to kill external MRSA cells, then host cells were treated with either loratadine, compound 8, or an untreated (DMSO) control. Three T-25 flasks were used per treatment. At 24 hours post-infection, two to three flasks were harvested per treatment for bacterial cell lysates. An additional flask per treatment had its medium replaced with DMEM containing no drugs. These flasks were imaged daily for morphological changes.

### RT-qPCR

Total RNA was purified from internalized bacterial lysates using lysostaphin and the QIAGEN RNeasy Kit with QIAshredder spin columns according to manufacturer’s instructions. Genomic DNA was eliminated with the Invitrogen TURBO DNA-free Kit. Reverse transcription was performed on 100ng of total RNA using New England Biolabs LunaScript RT SuperMix.

Quantitative PCR was performed using New England Biolabs Luna Universal qPCR Master Mix. Technical duplicates were performed, and negative controls for genomic DNA contamination (-RT) and reagent contamination (no template controls) were included. Results are shown from biological triplicate experiments. Each primer pair had its amplification efficiency experimentally determined. Primer details are found in Table S45. The comparative Ct method was used for relative quantification, with 16S rRNA as a reference gene.

### Morphological measurements

EA.hy926 and HeLa cells were photographed using an inverted light microscope, taking brightfield images with a 10X objective at the times indicated in the figures. NIH ImageJ software was used to measure roundness of 10 cells per field in each treatment. Results are averages from a representative experiment. GraphPad Prism v10.3.1 was used to analyze data with 2-way ANOVAs that were between measurements with no repeated measures. Drug treatment served as the column factor and time served as the row factor.

Interaction effects were included. Adjusted p-values were generated through multiple hypothesis testing using Tukey’s test.

## Supporting information

Dillon and Marshall et al Supporting Information

Dillon and Marshall et al Supporting Tables S35-37

Dillon and Marshall et al Supporting Tables S32-34

Dillon and Marshall et al Supporting Table S31

## Acknowledgements

Funding was provided by the National Institutes of Health (R15GM134503) to H.B.M. and M.S.B., the Camille and Henry Dreyfus Foundation to H.B.M., and the High Point University Natural Sciences Fellows Program.

---

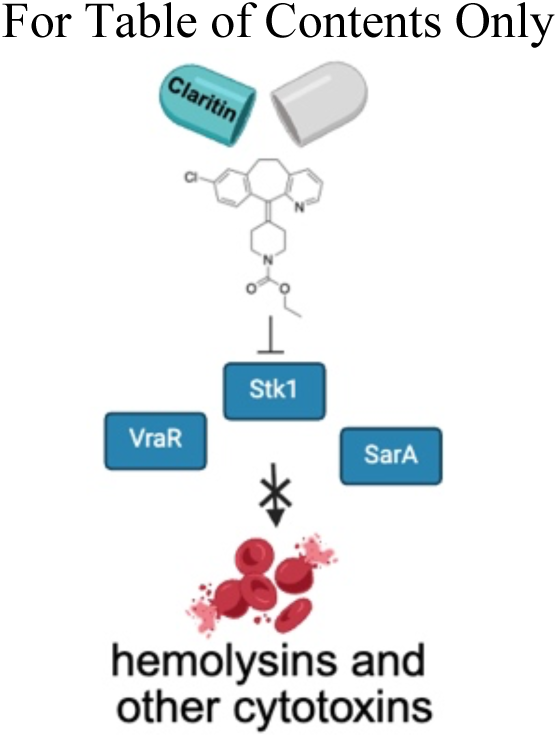

