## Supplementary material for "Loratadine decreases methicillin-resistant *Staphylococcus aureus* virulence through global hemolysin regulators *vraSR* and *agr*": Dillon and Marshall et al Supporting Information

**Table of Contents**

| **Supporting Tables** | **Page** |
|---|---|
| S1 - 2-way ANOVA results summarizing effects of loratadine-induced changes to hemolysis | 4 |
| S2 - 2-way ANOVA results analyzing effects of loratadine and stk1 on alpha hemolysis | 5 |
| S3 - 2-way ANOVA results analyzing effects of loratadine and stk1 on beta hemolysis | 5 |
| S4 - 2-way ANOVA results analyzing effects of loratadine and stp1 on alpha hemolysis | 5 |
| S5 - 2-way ANOVA results analyzing effects of loratadine and stp1 on beta hemolysis | 6 |
| S6 - 2-way ANOVA results summarizing effects of compound 8-induced changes to hemolysis | 6 |
| S7 - 2-way ANOVA results analyzing effects of loratadine and sarA on alpha hemolysis | 7 |
| S8 - 2-way ANOVA results analyzing effects of loratadine and sarZ on alpha hemolysis | 7 |
| S9 - 2-way ANOVA results analyzing effects of loratadine and sarA on beta hemolysis | 7 |
| S10 - 2-way ANOVA results analyzing effects of loratadine and sarZ on beta hemolysis | 8 |
| S11 - 2-way ANOVA results analyzing effects of loratadine and sarS on alpha hemolysis | 8 |
| S12 - 2-way ANOVA results analyzing effects of loratadine and sarT on alpha hemolysis | 8 |
| S13 - 2-way ANOVA results analyzing effects of loratadine and sarU on alpha hemolysis | 9 |
| S14 - 2-way ANOVA results analyzing effects of loratadine and sarV on alpha hemolysis | 9 |
| S15 - 2-way ANOVA results analyzing effects of loratadine and sarX on alpha hemolysis | 9 |
| S16 - 2-way ANOVA results analyzing effects of loratadine and sarS on beta hemolysis | 10 |
| S17 - 2-way ANOVA results analyzing effects of loratadine and sarT on beta hemolysis | 10 |
| S18 - 2-way ANOVA results analyzing effects of loratadine and sarU on beta hemolysis | 10 |
| S19 - 2-way ANOVA results analyzing effects of loratadine and sarV on beta hemolysis | 11 |
| S20 - 2-way ANOVA results analyzing effects of loratadine and sarX on beta hemolysis | 11 |
| S21 - 2-way ANOVA results analyzing effects of loratadine and saeR on alpha hemolysis | 11 |

**Supporting Tables** **Page**

S22 - 2-way ANOVA results analyzing effects of loratadine and saeS on alpha hemolysis 12
S23 - 2-way ANOVA results analyzing effects of loratadine and saeR on beta hemolysis 12
S24 - 2-way ANOVA results analyzing effects of loratadine and saeS on beta hemolysis 12
S25 - 2-way ANOVA results analyzing effects of loratadine and arlR on alpha hemolysis 13
S26 - 2-way ANOVA results analyzing effects of loratadine and arlS on alpha hemolysis 13
S27 - 2-way ANOVA results analyzing effects of loratadine and arlR on beta hemolysis 13
S28 - 2-way ANOVA results analyzing effects of loratadine and arlS on beta hemolysis 14
S29 - 2-way ANOVA results analyzing effects of loratadine and codY on alpha hemolysis 14
S30 - 2-way ANOVA results analyzing effects of loratadine and codY on beta hemolysis 14
S31-Complete DEG data sets for loratadine available in an Excel workbook
S32-34 – Loratadine GO Enrichment results available in an Excel workbook
S35-37 – Loratadine KEGG Enrichment results available in an Excel workbook
S38 - 2-way ANOVA results analyzing effects of loratadine and time on infected EA.hy926 cell roundness 15
S39 - 2-way ANOVA results analyzing effects of loratadine and time on uninfected EA.hy926 cell roundness 15
S40 - 2-way ANOVA results analyzing effects of loratadine and time on uninfected HeLa cell roundness 15
S41- Human cell line infection model RT-qPCR results with low/no amplification 16
S42 – Additional quality control details of RNA samples 16
S43 – RNA-seq read quality control 17
S44 – Mapped reads summary 17
S45 - qPCR primer information 18

**Supporting Figures** **Page**

S1 - Quantitative hemolysis assays analyzing effects of compound 8 and sarA and sarZ 18
S2 - Quantitative hemolysis assays analyzing effects of compound 8 and other sar genes 19
S3 - Quantitative hemolysis assays analyzing effects of compound 8 and saeR and saeS 19
S4 - Quantitative hemolysis assays analyzing effects of compound 8 and arlR and arlS 20
S5 - Quantitative hemolysis assays analyzing effects of compound 8 and codY 20
S6 – RT-qPCR validation of loratadine modulating multiple hemolysin genes in vitro 21
S7 - Temporary rounding of host cells treated with loratadine 22
S8 - Cytotoxicity of compound 8 and hemolysin mRNA levels in the human cell line infection model 23
S9 - Agarose gel electrophoresis of RT-qPCR products 24

**Table S1: 2-way ANOVA results summarizing effects of loratadine-induced changes to hemolysis.** The gene disruption and loratadine treatment were the column and row factors, respectively in the analysis. NS = not statistically significant in this analysis (p>0.05), NA = not applicable given that there is a significant interaction effect making column and row effects difficult to interpret. Green highlighted cells indicate mutant strains that produced statistically significant interaction effects in either alpha or beta hemolysin activity.

| | **Rabbit blood (alpha hemolysin)** | | | **Sheep blood (beta hemolysin)** | | |
|---|---|---|---|---|---|---|
| **Tn Mutant Strain** | **Gene Disruption Effect** | **Loratadine Effect** | **Interaction Effect** | **Gene Disruption Effect** | **Loratadine Effect** | **Interaction Effect** |
| Tn::stk1 | NA | NA | p=0.0379 | NS | p=0.0003 | NS |
| Tn::stp1 | NS | p=0.0341 | NS | NS | p=0.0135 | NS |
| Tn::sarA | NS | p=0.0547 | NS | NA | NA | p=0.0072 |
| Tn::sarZ | NS | p=0.0351 | NS | NS | p=0.0118 | NS |
| Tn::sarS | p=0.0004 | p=0.0455 | NS | NA | NA | p=0.0470 |
| Tn::sarT | NS | p=0.0452 | NS | NA | NA | p=0.0470 |
| Tn::sarU | NA | NA | p=0.0060 | NS | p=0.0015 | NS |
| Tn::sarV | NS | NS | NS | NS | p=0.0066 | NS |
| Tn::sarX | NA | NA | p=0.0037 | NS | p=0.0036 | NS |
| Tn::saeR | p<0.0001 | p=0.0011 | NS | NS | p=0.0009 | NS |
| Tn::saeS | p<0.0001 | p=0.0485 | NS | p=0.0404 | p=0.0022 | NS |
| Tn::arlR | NS | p=0.0484 | NS | NS | p=0.001 | NS |
| Tn::arlS | NA | NA | p=0.0233 | NS | p=0.003 | NS |
| Tn::codY | NA | NA | p=0.0385 | NS | p=0.001 | NS |

**Table S2: 2-way ANOVA results analyzing effects of loratadine and stk1 on alpha hemolysis.** The gene disruption and loratadine treatment were the column and row factors, respectively in the analysis. ns = not statistically significant in this analysis (p>0.05).

| **Source of Variation** | % total variation | P value | P value summary | Significant? | |
|---|---|---|---|---|---|
| Interaction | 15.32 | 0.0379 | * | Yes | |
| Loratadine | 47.21 | 0.0024 | ** | Yes | |
| Gene disruption of stk1 | 17.59 | 0.0288 | * | Yes | |
| **ANOVA table** | SS | DF | MS | F (DFn, DFd) | P value |
| Interaction | 1731 | 1 | 1731 | F (1, 8) = 6.164 | P=0.0379 |
| Loratadine | 5338 | 1 | 5338 | F (1, 8) = 19.00 | P=0.0024 |
| Gene disruption of stk1 | 1989 | 1 | 1989 | F (1, 8) = 7.081 | P=0.0288 |
| Residual | 2247 | 8 | 280.9 | | |

**Table S3: 2-way ANOVA results analyzing effects of loratadine and stp1 on alpha hemolysis.** The gene disruption and loratadine treatment were the column and row factors, respectively in the analysis. ns = not statistically significant in this analysis (p>0.05).

| **Source of Variation** | % total variation | P value | P value summary | Significant? | |
|---|---|---|---|---|---|
| Interaction | 11.17 | 0.1851 | ns | No | |
| Loratadine | 34.57 | 0.0341 | * | Yes | |
| Gene disruption of stp1 | 11.79 | 0.1745 | ns | No | |
| **ANOVA table** | SS | DF | MS | F (DFn, DFd) | P value |
| Interaction | 129.9 | 1 | 129.9 | F (1, 8) = 2.103 | P=0.1851 |
| Loratadine | 402.1 | 1 | 402.1 | F (1, 8) = 6.510 | P=0.0341 |
| Gene disruption of stp1 | 137.1 | 1 | 137.1 | F (1, 8) = 2.220 | P=0.1745 |
| Residual | 494.1 | 8 | 61.76 | | |

**Table S4: 2-way ANOVA results analyzing effects of loratadine and stk1 on beta hemolysis.** The gene disruption and loratadine treatment were the column and row factors, respectively in the analysis. ns = not statistically significant in this analysis (p>0.05).

| **Source of Variation** | % total variation | P value | P value summary | Significant? | |
|---|---|---|---|---|---|
| Interaction | 1.181 | 0.4744 | ns | No | |
| Loratadine | 77.87 | 0.0003 | *** | Yes | |
| Gene disruption of stk1 | 4.172 | 0.1961 | ns | No | |
| **ANOVA table** | SS | DF | MS | F (DFn, DFd) | P value |
| Interaction | 44.42 | 1 | 44.42 | F (1, 8) = 0.5634 | P=0.4744 |
| Loratadine | 2928 | 1 | 2928 | F (1, 8) = 37.14 | P=0.0003 |
| Gene disruption of stk1 | 156.9 | 1 | 156.9 | F (1, 8) = 1.990 | P=0.1961 |
| Residual | 630.8 | 8 | 78.85 | | |

**Table S5: 2-way ANOVA results analyzing effects of loratadine and stp1 on beta hemolysis.** The gene disruption and loratadine treatment were the column and row factors, respectively in the analysis. ns = not statistically significant in this analysis (p>0.05).

| **Source of Variation** | % total variation | P value | P value summary | Significant? | |
|---|---|---|---|---|---|
| Interaction | 8.644 | 0.2052 | ns | No | |
| Loratadine | 45.21 | 0.0135 | * | Yes | |
| Gene disruption stp1 | 9.781 | 0.1806 | ns | No | |
| **ANOVA table** | **SS** | **DF** | **MS** | **F (DFn, DFd)** | **P value** |
| Interaction | 341.9 | 1 | 341.9 | F (1, 8) = 1.902 | P=0.2052 |
| Loratadine | 1788 | 1 | 1788 | F (1, 8) = 9.946 | P=0.0135 |
| Gene disruption stp1 | 386.9 | 1 | 386.9 | F (1, 8) = 2.152 | P=0.1806 |
| Residual | 1438 | 8 | 179.8 | | |

**Table S6: 2-way ANOVA summary results analyzing effects of compound 8-induced changes to hemolysis.** The gene disruption and compound 8 treatment were the column and row factors, respectively in the analysis. NS = not statistically significant in this analysis (p>0.05).

| | **Rabbit blood (alpha hemolysin)** | | | **Sheep blood (beta hemolysin)** | | |
|---|---|---|---|---|---|---|
| **Transposon Mutant Strain** | **Gene Disruption Effect** | **Compound 8 Effect** | **Interaction Effect** | **Gene Disruption Effect** | **Compound 8 Effect** | **Interaction Effect** |
| Tn::stk1 | p=0.0353 | NS | NS | NS | NS | NS |
| Tn::stp1 | NS | NS | NS | NS | NS | NS |
| Tn::sarA | p=0.0156 | NS | NS | p=0.0008 | NS | NS |
| Tn::sarZ | NS | NS | NS | NS | NS | NS |
| Tn::sarS | p=0.0013 | NS | NS | p=0.157 | NS | NS |
| Tn::sarT | NS | NS | NS | p=0.0145 | NS | NS |
| Tn::sarU | p=0.0001 | NS | NS | p=0.0419 | NS | NS |
| Tn::sarV | p<0.0001 | NS | NS | NS | NS | NS |
| Tn::sarX | p=0.0095 | NS | NS | NS | NS | NS |
| Tn::saeR | p<0.0001 | NS | NS | NS | NS | NS |
| Tn::saeS | p<0.0001 | NS | NS | p=0.0412 | NS | NS |
| Tn::arlR | NS | NS | NS | NS | NS | NS |
| Tn::arlS | p=0.0022 | NS | NS | NS | NS | NS |
| Tn::codY | p<0.0001 | NS | NS | NS | NS | NS |

**Table S7: 2-way ANOVA results analyzing effects of loratadine and *sarA* on alpha hemolysis.** The gene disruption and loratadine treatment were the column and row factors, respectively in the analysis. ns = not statistically significant in this analysis (p>0.05).

| **Source of Variation** | **% total variation** | **P value** | **P value summary** | **Significant?** | |
|---|---|---|---|---|---|
| Interaction | 15.99 | 0.1524 | ns | No | |
| Loratadine | 32.32 | 0.0547 | ns | No | |
| Gene disruption sarA | 0.5486 | 0.7770 | ns | No | |
| **ANOVA table** | **SS** | **DF** | **MS** | **F (DFn, DFd)** | **P value** |
| Interaction | 168.6 | 1 | 168.6 | F (1, 8) = 2.501 | P=0.1524 |
| Loratadine | 340.8 | 1 | 340.8 | F (1, 8) = 5.055 | P=0.0547 |
| Gene disruption sarA | 5.786 | 1 | 5.786 | F (1, 8) = 0.08581 | P=0.7770 |
| Residual | 539.4 | 8 | 67.42 | | |

**Table S8: 2-way ANOVA results analyzing effects of loratadine and *sarZ* on alpha hemolysis.** The gene disruption and loratadine treatment were the column and row factors, respectively in the analysis. ns = not statistically significant in this analysis (p>0.05).

| **Source of Variation** | **% total variation** | **P value** | **P value summary** | **Significant** | |
|---|---|---|---|---|---|
| Interaction | 0.3719 | 0.8221 | ns | No | |
| Loratadine | 44.19 | 0.0351 | * | Yes | |
| Gene disruption | 0.3484 | 0.8277 | ns | No | |
| **ANOVA table** | **SS** | **DF** | **MS** | **F (DFn,** | **P value** |
| Interaction | 6.983 | 1 | 6.983 | F (1, 8) = | P=0.8221 |
| Loratadine | 829.7 | 1 | 829.7 | F (1, 8) = | P=0.0351 |
| Gene disruption | 6.542 | 1 | 6.542 | F (1, 8) = | P=0.8277 |
| Residual | 1034 | 8 | 129.3 | | |

**Table S9: 2-way ANOVA results analyzing effects of loratadine and *sarA* on beta hemolysis.** The gene disruption and loratadine treatment were the column and row factors, respectively in the analysis. ns = not statistically significant in this analysis (p>0.05).

| **Source of Variation** | **% total variation** | **P value** | **P value summary** | **Significant?** | |
|---|---|---|---|---|---|
| Interaction | 23.74 | 0.0072 | ** | Yes | |
| Loratadine | 21.39 | 0.0094 | ** | Yes | |
| Gene disruption | 40.03 | 0.0017 | ** | Yes | |
| **ANOVA table** | **SS** | **DF** | **MS** | **F (DFn, DFd)** | **P value** |
| Interaction | 972.1 | 1 | 972.1 | F (1, 8) = 12.79 | P=0.0072 |
| Loratadine | 876.0 | 1 | 876.0 | F (1, 8) = 11.53 | P=0.0094 |
| Gene disruption | 1639 | 1 | 1639 | F (1, 8) = 21.57 | P=0.0017 |
| Residual | 608.1 | 8 | 76.01 | | |

**Table S10: 2-way ANOVA results analyzing effects of loratadine and *sarZ* on beta hemolysis.** The gene disruption and loratadine treatment were the column and row factors, respectively in the analysis. ns = not statistically significant in this analysis (p>0.05).

| **Source of Variation** | **% total variation** | **P value** | **P value summary** | **Significant?** | |
|---|---|---|---|---|---|
| Interaction | 3.245 | 0.4524 | ns | No | |
| Loratadine | 54.70 | 0.0118 | * | Yes | |
| Gene disruption | 0.4360 | 0.7796 | ns | No | |
| **ANOVA table** | **SS** | **DF** | **MS** | **F (DFn, DFd)** | **P value** |
| Interaction | 141.7 | 1 | 141.7 | F (1, 8) = 0.6237 | P=0.4524 |
| Loratadine | 2389 | 1 | 2389 | F (1, 8) = 10.51 | P=0.0118 |
| Gene disruption | 19.04 | 1 | 19.04 | F (1, 8) = 0.08382 | P=0.7796 |
| Residual | 1817 | 8 | 227.2 | | |

**Table S11: 2-way ANOVA results analyzing effects of loratadine and *sarS* on alpha hemolysis.** The gene disruption and loratadine treatment were the column and row factors, respectively in the analysis. ns = not statistically significant in this analysis (p>0.05).

| **Source of Variation** | **% total variation** | **P value** | **P value summary** | **Significant?** | |
|---|---|---|---|---|---|
| Interaction | 1.469 | 0.4212 | ns | No | |
| Loratadine | 11.45 | 0.0455 | * | Yes | |
| Gene disruption | 70.73 | 0.0004 | *** | Yes | |
| **ANOVA table** | **SS** | **DF** | **MS** | **F (DFn, DFd)** | **P value** |
| Interaction | 308.0 | 1 | 308.0 | F (1, 8) = 0.7187 | P=0.4212 |
| Loratadine | 2401 | 1 | 2401 | F (1, 8) = 5.602 | P=0.0455 |
| Gene disruption | 14827 | 1 | 14827 | F (1, 8) = 34.60 | P=0.0004 |
| Residual | 3428 | 8 | 428.5 | | |

**Table S12: 2-way ANOVA results analyzing effects of loratadine and *sarT* on alpha hemolysis.** The gene disruption and loratadine treatment were the column and row factors, respectively in the analysis. ns = not statistically significant in this analysis (p>0.05).

| **Source of Variation** | **% total variation** | **P value** | **P value summary** | **Significant?** | |
|---|---|---|---|---|---|
| Interaction | 0.0001845 | 0.9962 | ns | No | |
| Loratadine | 45.25 | 0.0452 | * | Yes | |
| Gene disruption sarT | 0.02205 | 0.9587 | ns | No | |
| **ANOVA table** | **SS** | **DF** | **MS** | **F (DFn, DFd)** | **P value** |
| Interaction | 0.003598 | 1 | 0.003598 | F (1, 7) = 2.413e-005 | P=0.9962 |
| Loratadine | 882.6 | 1 | 882.6 | F (1, 7) = 5.920 | P=0.0452 |
| Gene disruption sarT | 0.4300 | 1 | 0.4300 | F (1, 7) = 0.002884 | P=0.9587 |
| Residual | 1044 | 7 | 149.1 | | |

**Table S13: 2-way ANOVA results analyzing effects of loratadine and *sarU* on alpha hemolysis.** The gene disruption and loratadine treatment were the column and row factors, respectively in the analysis. ns = not statistically significant in this analysis (p>0.05).

| **Source of Variation** | **% total variation** | **P value** | **P value summary** | **Significant?** | |
|---|---|---|---|---|---|
| Interaction | 10.66 | 0.0060 | ** | Yes | |
| Loratadine | 31.40 | 0.0002 | *** | Yes | |
| Gene disruption | 51.74 | <0.0001 | **** | Yes | |
| **ANOVA table** | **SS** | **DF** | **MS** | **F (DFn, DFd)** | **P value** |
| Interaction | 1928 | 1 | 1928 | F (1, 8) = 13.75 | P=0.0060 |
| Loratadine | 5679 | 1 | 5679 | F (1, 8) = 40.50 | P=0.0002 |
| Gene disruption | 9360 | 1 | 9360 | F (1, 8) = 66.74 | P<0.0001 |
| Residual | 1122 | 8 | 140.2 | | |

**Table S14: 2-way ANOVA results analyzing effects of loratadine and *sarV* on alpha hemolysis.** The gene disruption and loratadine treatment were the column and row factors, respectively in the analysis. ns = not statistically significant in this analysis (p>0.05).

| **Source of Variation** | **% total variation** | **P value** | **P value** | **Significant?** | |
|---|---|---|---|---|---|
| Interaction | 1.641 | 0.6638 | ns | No | |
| Loratadine | 33.67 | 0.0752 | ns | No | |
| Gene disruption | 0.2302 | 0.8700 | ns | No | |
| **ANOVA table** | **SS** | **DF** | **MS** | **F (DFn, DFd)** | **P value** |
| Interaction | 79.37 | 1 | 79.37 | F (1, 8) = 0.2037 | P=0.6638 |
| Loratadine | 1629 | 1 | 1629 | F (1, 8) = 4.179 | P=0.0752 |
| Gene disruption | 11.13 | 1 | 11.13 | F (1, 8) = 0.02857 | P=0.8700 |
| Residual | 3118 | 8 | 389.7 | | |

**Table S15: 2-way ANOVA results analyzing effects of loratadine and *sarX* on alpha hemolysis.** The gene disruption and loratadine treatment were the column and row factors, respectively in the analysis. ns = not statistically significant in this analysis (p>0.05).

| **Source of Variation** | **% total variation** | **P value** | **P value** | **Significant?** | |
|---|---|---|---|---|---|
| Interaction | 13.85 | 0.0037 | ** | Yes | |
| Loratadine | 36.73 | 0.0002 | *** | Yes | |
| Gene disruption | 42.67 | 0.0001 | *** | Yes | |
| **ANOVA table** | **SS** | **DF** | **MS** | **F (DFn, DFd)** | **P value** |
| Interaction | 2502 | 1 | 2502 | F (1, 8) = 16.41 | P=0.0037 |
| Loratadine | 6637 | 1 | 6637 | F (1, 8) = 43.52 | P=0.0002 |
| Gene disruption | 7710 | 1 | 7710 | F (1, 8) = 50.56 | P=0.0001 |
| Residual | 1220 | 8 | 152.5 | | |

**Table S16: 2-way ANOVA results analyzing effects of loratadine and *sarS* on beta hemolysis.** The gene disruption and loratadine treatment were the column and row factors, respectively in the analysis. ns = not statistically significant in this analysis (p>0.05).

| **Source of Variation** | **% total variation** | **P value** | **P value** | **Significant?** | |
|---|---|---|---|---|---|
| Interaction | 20.59 | 0.0313 | * | Yes | |
| Loratadine | 36.98 | 0.0082 | ** | Yes | |
| Gene disruption | 18.18 | 0.0400 | * | Yes | |
| **ANOVA table** | **SS** | **DF** | **MS** | **F (DFn, DFd)** | **P value** |
| Interaction | 674.5 | 1 | 674.5 | F (1, 8) = 6.793 | P=0.0313 |
| Loratadine | 1211 | 1 | 1211 | F (1, 8) = 12.20 | P=0.0082 |
| Gene disruption | 595.4 | 1 | 595.4 | F (1, 8) = 5.997 | P=0.0400 |
| Residual | 794.3 | 8 | 99.29 | | |

**Table S17: 2-way ANOVA results analyzing effects of loratadine and *sarT* on beta hemolysis.** The gene disruption and loratadine treatment were the column and row factors, respectively in the analysis. ns = not statistically significant in this analysis (p>0.05).

| **Source of Variation** | **% total variation** | **P value** | **P value summary** | **Significant?** | |
|---|---|---|---|---|---|
| Interaction | 17.22 | 0.0470 | * | Yes | |
| Loratadine | 41.20 | 0.0067 | ** | Yes | |
| Gene disruption | 16.56 | 0.0504 | ns | No | |
| **ANOVA table** | **SS** | **DF** | **MS** | **F (DFn, DFd)** | **P value** |
| Interaction | 569.4 | 1 | 569.4 | F (1, 8) = 5.503 | P=0.0470 |
| Loratadine | 1363 | 1 | 1363 | F (1, 8) = 13.17 | P=0.0067 |
| Gene disruption | 547.7 | 1 | 547.7 | F (1, 8) = 5.294 | P=0.0504 |
| Residual | 827.7 | 8 | 103.5 | | |

**Table S18: 2-way ANOVA results analyzing effects of loratadine and *sarU* on beta hemolysis.** The gene disruption and loratadine treatment were the column and row factors, respectively in the analysis. ns = not statistically significant in this analysis (p>0.05).

| **Source of Variation** | **% total variation** | **P value** | **P value** | **Significant?** | |
|---|---|---|---|---|---|
| Interaction | 8.184 | 0.1160 | ns | No | |
| Loratadine | 58.65 | 0.0015 | ** | Yes | |
| Gene disruption | 12.09 | 0.0645 | ns | No | |
| **ANOVA table** | **SS** | **DF** | **MS** | **F (DFn, DFd)** | **P value** |
| Interaction | 273.2 | 1 | 273.2 | F (1, 8) = 3.107 | P=0.1160 |
| Loratadine | 1958 | 1 | 1958 | F (1, 8) = 22.27 | P=0.0015 |
| Gene disruption | 403.6 | 1 | 403.6 | F (1, 8) = 4.590 | P=0.0645 |
| Residual | 703.5 | 8 | 87.93 | | |

**Table S19: 2-way ANOVA results analyzing effects of loratadine and *sarV* on beta hemolysis.** The gene disruption and loratadine treatment were the column and row factors, respectively in the analysis. ns = not statistically significant in this analysis (p>0.05).

| Source of Variation | % total variation | P value | P value | Significant? | |
|---|---|---|---|---|---|
| Interaction | 16.99 | 0.0633 | ns | No | |
| Loratadine | 48.37 | 0.0066 | ** | Yes | |
| Gene disruption | 5.358 | 0.2608 | ns | No | |
| **ANOVA table** | **SS** | **DF** | **MS** | **F (DFn, DFd)** | **P value** |
| Interaction | 511.6 | 1 | 511.6 | F (1, 8) = 4.644 | P=0.0633 |
| Loratadine | 1456 | 1 | 1456 | F (1, 8) = 13.22 | P=0.0066 |
| Gene disruption | 161.3 | 1 | 161.3 | F (1, 8) = 1.464 | P=0.2608 |
| Residual | 881.4 | 8 | 110.2 | | |

**Table S20: 2-way ANOVA results analyzing effects of loratadine and *sarX* on beta hemolysis.** The gene disruption and loratadine treatment were the column and row factors, respectively in the analysis. ns = not statistically significant in this analysis (p>0.05).

| Source of Variation | % total variation | P value | P value | Significant? | |
|---|---|---|---|---|---|
| Interaction | 1.499 | 0.5539 | ns | No | |
| Loratadine | 64.76 | 0.0036 | ** | Yes | |
| Gene disruption | 2.308 | 0.4655 | ns | No | |
| **ANOVA table** | **SS** | **DF** | **MS** | **F (DFn, DFd)** | **P value** |
| Interaction | 64.41 | 1 | 64.41 | F (1, 8) = 0.3816 | P=0.5539 |
| Loratadine | 2783 | 1 | 2783 | F (1, 8) = 16.48 | P=0.0036 |
| Gene disruption | 99.15 | 1 | 99.15 | F (1, 8) = 0.5873 | P=0.4655 |
| Residual | 1350 | 8 | 168.8 | | |

**Table S21: 2-way ANOVA results analyzing effects of loratadine and *saeR* on alpha hemolysis.** The gene disruption and loratadine treatment were the column and row factors, respectively in the analysis. ns = not statistically significant in this analysis (p>0.05).

| Source of Variation | % total variation | P value | P value | Significant? | |
|---|---|---|---|---|---|
| Interaction | 0.5033 | 0.2726 | ns | No | |
| Loratadine | 8.986 | 0.0011 | ** | Yes | |
| Gene disruption | 87.61 | <0.0001 | **** | Yes | |
| **ANOVA table** | **SS** | **DF** | **MS** | **F (DFn, DFd)** | **P value** |
| Interaction | 95.05 | 1 | 95.05 | F (1, 8) = 1.388 | P=0.2726 |
| Loratadine | 1697 | 1 | 1697 | F (1, 8) = 24.78 | P=0.0011 |
| Gene disruption | 16547 | 1 | 16547 | F (1, 8) = 241.6 | P<0.0001 |
| Residual | 547.9 | 8 | 68.49 | | |

**Table S22: 2-way ANOVA results analyzing effects of loratadine and *saeS* on alpha hemolysis.** The gene disruption and loratadine treatment were the column and row factors, respectively in the analysis. ns = not statistically significant in this analysis (p>0.05).

| Source of Variation | % total variation | P value | P value | Significant? | |
|---|---|---|---|---|---|
| Interaction | 0.7208 | 0.0956 | ns | No | |
| Loratadine | 1.093 | 0.0485 | * | Yes | |
| Gene disruption | 96.57 | <0.0001 | **** | Yes | |
| **ANOVA table** | **SS** | **DF** | **MS** | **F (DFn, DFd)** | **P value** |
| Interaction | 198.6 | 1 | 198.6 | F (1, 8) = 3.568 | P=0.0956 |
| Loratadine | 301.2 | 1 | 301.2 | F (1, 8) = 5.410 | P=0.0485 |
| Gene disruption | 26610 | 1 | 26610 | F (1, 8) = 478.0 | P<0.0001 |
| Residual | 445.3 | 8 | 55.67 | | |

**Table S23: 2-way ANOVA results analyzing effects of loratadine and *saeR* on beta hemolysis.** The gene disruption and loratadine treatment were the column and row factors, respectively in the analysis. ns = not statistically significant in this analysis (p>0.05).

| Source of Variation | % total variation | P value | P value | Significant? | |
|---|---|---|---|---|---|
| Interaction | 6.752 | 0.1407 | ns | No | |
| Loratadine | 66.22 | 0.0009 | *** | Yes | |
| Gene disruption | 6.822 | 0.1389 | ns | No | |
| **ANOVA table** | **SS** | **DF** | **MS** | **F (DFn, DFd)** | **P value** |
| Interaction | 216.4 | 1 | 216.4 | F (1, 8) = 2.673 | P=0.1407 |
| Loratadine | 2122 | 1 | 2122 | F (1, 8) = 26.22 | P=0.0009 |
| Gene disruption | 218.6 | 1 | 218.6 | F (1, 8) = 2.701 | P=0.1389 |
| Residual | 647.6 | 8 | 80.95 | | |

**Table S24: 2-way ANOVA results analyzing effects of loratadine and *saeS* on beta hemolysis.** The gene disruption and loratadine treatment were the column and row factors, respectively in the analysis. ns = not statistically significant in this analysis (p>0.05).

| Source of Variation | % total variation | P value | P value summary | Significant? | |
|---|---|---|---|---|---|
| Interaction | 13.53 | 0.0510 | ns | No | |
| Loratadine | 50.53 | 0.0022 | ** | Yes | |
| Gene disruption | 15.35 | 0.0404 | * | Yes | |
| **ANOVA table** | **SS** | **DF** | **MS** | **F (DFn, DFd)** | **P value** |
| Interaction | 429.6 | 1 | 429.6 | F (1, 8) = 5.259 | P=0.0510 |
| Loratadine | 1604 | 1 | 1604 | F (1, 8) = 19.63 | P=0.0022 |
| Gene disruption | 487.3 | 1 | 487.3 | F (1, 8) = 5.965 | P=0.0404 |
| Residual | 653.5 | 8 | 81.69 | | |

**Table S25: 2-way ANOVA results analyzing effects of loratadine and *arlR* on alpha hemolysis.** The gene disruption and loratadine treatment were the column and row factors, respectively in the analysis. ns = not statistically significant in this analysis (p>0.05).

| **Source of Variation** | **% total variation** | **P value** | **P value** | **Significant?** | |
|---|---|---|---|---|---|
| Interaction | 2.871 | 0.5385 | ns | No | |
| Loratadine | 37.63 | 0.0484 | * | Yes | |
| Gene disruption | 3.867 | 0.4772 | ns | No | |
| **ANOVA table** | **SS** | **DF** | **MS** | **F (DFn, DFd)** | **P value** |
| Interaction | 46.32 | 1 | 46.32 | F (1, 8) = 0.4129 | P=0.5385 |
| Loratadine | 607.2 | 1 | 607.2 | F (1, 8) = 5.412 | P=0.0484 |
| Gene disruption | 62.39 | 1 | 62.39 | F (1, 8) = 0.5561 | P=0.4772 |
| Residual | 897.5 | 8 | 112.2 | | |

**Table S26: 2-way ANOVA results analyzing effects of loratadine and *arlS* on alpha hemolysis.** The gene disruption and loratadine treatment were the column and row factors, respectively in the analysis. ns = not statistically significant in this analysis (p>0.05).

| **Source of Variation** | **% total variation** | **P value** | **P value** | **Significant?** | |
|---|---|---|---|---|---|
| Interaction | 10.62 | 0.0233 | * | Yes | |
| Loratadine | 41.17 | 0.0006 | *** | Yes | |
| Gene disruption | 37.35 | 0.0008 | *** | Yes | |
| **ANOVA table** | **SS** | **DF** | **MS** | **F (DFn, DFd)** | **P value** |
| Interaction | 1053 | 1 | 1053 | F (1, 8) = 7.828 | P=0.0233 |
| Loratadine | 4084 | 1 | 4084 | F (1, 8) = 30.34 | P=0.0006 |
| Gene disruption | 3705 | 1 | 3705 | F (1, 8) = 27.53 | P=0.0008 |
| Residual | 1077 | 8 | 134.6 | | |

**Table S27: 2-way ANOVA results analyzing effects of loratadine and *arlR* on beta hemolysis.** The gene disruption and loratadine treatment were the column and row factors, respectively in the analysis. ns = not statistically significant in this analysis (p>0.05).

| **Source of Variation** | **% total variation** | **P value** | **P value** | **Significant?** | |
|---|---|---|---|---|---|
| Interaction | 4.046 | 0.2683 | ns | No | |
| Loratadine | 71.66 | 0.0010 | ** | Yes | |
| Gene disruption | 1.422 | 0.5007 | ns | No | |
| **ANOVA table** | **SS** | **DF** | **MS** | **F (DFn, DFd)** | **P value** |
| Interaction | 136.2 | 1 | 136.2 | F (1, 8) = 1.415 | P=0.2683 |
| Loratadine | 2411 | 1 | 2411 | F (1, 8) = 25.06 | P=0.0010 |
| Gene disruption | 47.84 | 1 | 47.84 | F (1, 8) = 0.4972 | P=0.5007 |
| Residual | 769.8 | 8 | 96.22 | | |

**Table S28: 2-way ANOVA results analyzing effects of loratadine and *arlS* on beta hemolysis.** The gene disruption and loratadine treatment were the column and row factors, respectively in the analysis. ns = not statistically significant in this analysis (p>0.05).

| **Source of Variation** | **% total variation** | **P value** | **P value** | **Significant?** | |
|---|---|---|---|---|---|
| Interaction | 0.1525 | 0.8032 | ns | No | |
| Loratadine | 81.42 | 0.0003 | *** | Yes | |
| Gene disruption | 0.04945 | 0.8870 | ns | No | |
| **ANOVA table** | **SS** | **DF** | **MS** | **F (DFn, DFd)** | **P value** |
| Interaction | 6.356 | 1 | 6.356 | F (1, 8) = 0.06636 | P=0.8032 |
| Loratadine | 3394 | 1 | 3394 | F (1, 8) = 35.43 | P=0.0003 |
| Gene disruption | 2.061 | 1 | 2.061 | F (1, 8) = 0.02152 | P=0.8870 |
| Residual | 766.3 | 8 | 95.78 | | |

**Table S29: 2-way ANOVA results analyzing effects of loratadine and *codY* on alpha hemolysis.** The gene disruption and loratadine treatment were the column and row factors, respectively in the analysis. ns = not statistically significant in this analysis (p>0.05).

| **Source of Variation** | **% total variation** | **P value** | **P value** | **Significant?** | |
|---|---|---|---|---|---|
| Interaction | 7.728 | 0.0385 | * | Yes | |
| Loratadine | 33.57 | 0.0009 | *** | Yes | |
| Gene disruption | 48.59 | 0.0003 | *** | Yes | |
| **ANOVA table** | **SS** | **DF** | **MS** | **F (DFn, DFd)** | **P value** |
| Interaction | 841.1 | 1 | 841.1 | F (1, 8) = 6.119 | P=0.0385 |
| Loratadine | 3654 | 1 | 3654 | F (1, 8) = 26.58 | P=0.0009 |
| Gene disruption | 5289 | 1 | 5289 | F (1, 8) = 38.48 | P=0.0003 |
| Residual | 1100 | 8 | 137.5 | | |

**Table S30: 2-way ANOVA results analyzing effects of loratadine and *codY* on beta hemolysis.** The gene disruption and loratadine treatment were the column and row factors, respectively in the analysis. ns = not statistically significant in this analysis (p>0.05).

| **Source of Variation** | **% total variation** | **P value** | **P value** | **Significant?** | |
|---|---|---|---|---|---|
| Interaction | 1.525 | 0.4973 | ns | No | |
| Loratadine | 74.19 | 0.0011 | ** | Yes | |
| Gene disruption | 0.1444 | 0.8323 | ns | No | |
| **ANOVA table** | **SS** | **DF** | **MS** | **F (DFn, DFd)** | **P value** |
| Interaction | 58.09 | 1 | 58.09 | F (1, 8) = 0.5055 | P=0.4973 |
| Loratadine | 2825 | 1 | 2825 | F (1, 8) = 24.59 | P=0.0011 |
| Gene disruption | 5.499 | 1 | 5.499 | F (1, 8) = 0.04785 | P=0.8323 |
| Residual | 919.3 | 8 | 114.9 | | |

**Table S38: 2-way ANOVA results analyzing effects of loratadine and time on infected EA.hy926 cell roundness.** The time and loratadine treatment were the column and row factors, respectively in the analysis. ns = not statistically significant in this analysis (p>0.05).

| Source of Variation | % total variation | P value | P value | Significant? | |
|---|---|---|---|---|---|
| Interaction | 10.03 | 0.0350 | * | Yes | |
| Time | 4.932 | 0.0748 | ns | No | |
| Drug Treatment | 10.42 | 0.0050 | ** | Yes | |
| **ANOVA table** | **SS** | **DF** | **MS** | **F (DFn, DFd)** | **P value** |
| Interaction | 0.4571 | 4 | 0.1143 | F (4, 81) = 2.72 | P=0.0350 |
| Time | 0.2247 | 2 | 0.1124 | F (2, 81) = 2.67 | P=0.0748 |
| Drug Treatment | 0.4747 | 2 | 0.2374 | F (2, 81) = 5.65 | P=0.0050 |
| Residual | 3.400 | 81 | 0.04197 | | |

**Table S39: 2-way ANOVA results analyzing effects of loratadine and time on uninfected EA.hy926 cell roundness.** The time and loratadine treatment were the column and row factors, respectively in the analysis. ns = not statistically significant in this analysis (p>0.05).

| Source of Variation | % total variation | P value | P value summary | Significant? | |
|---|---|---|---|---|---|
| Interaction | 3.127 | 0.3487 | ns | No | |
| Time | 26.67 | <0.0001 | **** | Yes | |
| Drug Treatment | 14.11 | 0.0001 | *** | Yes | |
| **ANOVA table** | **SS** | **DF** | **MS** | **F (DFn, DFd)** | **P value** |
| Interaction | 0.1379 | 4 | 0.03448 | F (4, 80) = 1.129 | P=0.3487 |
| Time | 1.176 | 2 | 0.5881 | F (2, 80) = 19.26 | P<0.0001 |
| Drug Treatment | 0.6223 | 2 | 0.3112 | F (2, 80) = 10.19 | P=0.0001 |
| Residual | 2.443 | 80 | 0.03054 | | |

**Table S40: 2-way ANOVA results analyzing effects of loratadine and time on uninfected HeLa cell roundness.** The time and loratadine treatment were the column and row factors, respectively in the analysis. ns = not statistically significant in this analysis (p>0.05).

| Source of Variation | % total variation | P value | P value summary | Significant? | |
|---|---|---|---|---|---|
| Interaction | 11.45 | 0.0088 | ** | Yes | |
| Time | 7.983 | 0.0084 | ** | Yes | |
| Drug Treatment | 16.91 | <0.0001 | **** | Yes | |
| **ANOVA table** | **SS** | **DF** | **MS** | **F (DFn, DFd)** | **P value** |
| Interaction | 0.3160 | 4 | 0.07901 | F (4, 81) = 3.644 | P=0.0088 |
| Time | 0.2203 | 2 | 0.1101 | F (2, 81) = 5.079 | P=0.0084 |
| Drug Treatment | 0.4665 | 2 | 0.2332 | F (2, 81) = 10.76 | P<0.0001 |
| Residual | 1.756 | 81 | 0.02168 | | |

**Table S41: Human cell line infection model RT-qPCR results with low/no amplification.** Data shown are from a representative experiment with mean Ct values from duplicate reactions using untreated but infected cells. +RT = PCR reaction with cDNA as template, -RT = control PCR reaction that lacked reverse transcriptase in the cDNA synthesis step, NTC = no template control PCR reaction that lacked template and had only water and PCR mastermix. Red Ct values indicate that the Ct obtained in the +RT reaction was less than 5 Cts different from the –RT and NTC reactions. This indicates amplification of these genes is similar to background levels.

| Gene | WT JE2 infection | Tn::stk1 infection | Tn::stp1 infection | Tn::sarA infection |
|---|---|---|---|---|
| | **Mean Ct** | **Meant Ct** | **Meant Ct** | **Meant Ct** |
| **+RT** | | | | |
| 16S | 11.78793 | 25.29228 | 17.7547836 | 10.17088 |
| hla | 22.1402 | 32.09037 | 26.07072 | 20.2918625 |
| hlgA | 24.94802 | 31.7953 | 27.7055206 | 26.54224 |
| hlgC | 27.6679 | 28.31001 | 31.5830002 | 25.31952 |
| hld/RNAIII | 22.6522751 | 32.05889 | 27.99364 | 24.21673 |
| **-RT** | | | | |
| 16S | 30.59434 | 33.7104 | 34.16547 | 32.8718033 |
| hla | 28.657629 | 28.19838 | 27.77948 | 26.91212 |
| hlgA | 27.49573 | 32.69831 | 29.6728287 | 31.15741 |
| hlgC | 29.7959023 | 30.72838 | 32.3512535 | 25.31952 |
| hld/RNAIII | 27.39732 | 31.90166 | 32.8868256 | 34.2149353 |
| **NTC** | | | | |
| 16S | 29.67368 | 32.40137 | 34.86722 | 33.87514 |
| hla | 24.0518951 | 33.60343 | 27.81045 | 27.4429951 |
| hlgA | 28.35803 | 33.59329 | 30.29202 | 31.65088 |
| hlgC | 28.83661 | 31.24105 | 29.8993 | 31.47275 |
| hld/RNAIII | 31.9273 | 33.37765 | 31.96478 | 34.17091 |

**Table S42: Additional quality control details of RNA samples.** Un represents untreated sample, lor represents loratadine treated sample. Each biological replicate is labelled A, B, or C. RIN = RNA integrity number

| Sample Name | Concentration (ng/µL) | RIN |
|---|---|---|
| UnA | 64.8 | 6.4 |
| UnB | 279.2 | 7.8 |
| UnC | 202.0 | 7.7 |
| LorA | 43.2 | 5.8 |
| LorB | 171.2 | 7.1 |
| LorC | 76.4 | 7.7 |

**Table S43: RNA-seq reads quality control summary.** Un represents untreated sample, lor represents loratadine treated sample. Each biological replicate is labelled A, B, or C. Q20 and Q30 were calculated as the base number of Phred value > 20 or 30, respectively, divided by the total base value x 100%.

| Sample name | Raw reads | Clean reads | Raw bases | Clean bases | Error rate | Q20 | Q30 | GC content |
|---|---|---|---|---|---|---|---|---|
| UnA | 24119548 | 22181216 | 3.62G | 3.33G | 0.03 | 97.37 | 92.5 | 34.63 |
| LorA | 22706338 | 20878286 | 3.41G | 3.14G | 0.03 | 97.02 | 92.00 | 33.91 |
| UnB | 27424072 | 26201028 | 4.12G | 3.94G | 0.03 | 97.23 | 92.16 | 34.49 |
| LorB | 26122322 | 25020814 | 3.92G | 3.76G | 0.03 | 97.23 | 92.18 | 34.33 |
| UnC | 26598234 | 25372060 | 3.99G | 3.81G | 0.03 | 97.29 | 92.33 | 34.61 |
| LorC | 26059026 | 24610654 | 3.91G | 3.7G | 0.03 | 97.31 | 92.32 | 34.42 |

**Table S44: Mapped reads summary.** Un represents untreated sample, lor represents loratadine treated sample. Each biological replicate is labelled A, B, or C.

| **Sample name** | **UnA** | **UnB** | **UnC** | **LorA** | **LorB** | **LorC** |
|---|---|---|---|---|---|---|
| **Total reads** | 22181216 | 26201028 | 25372060 | 20878286 | 25020814 | 24610654 |
| **Total mapped** | 21817247 (98.36%) | 25740613 (98.24%) | 24961910 (98.38%) | 19796912 (94.82%) | 24449618 (97.72%) | 24117202 (97.99%) |
| **Multiple mapped** | 428988 (1.93%) | 527355 (2.01%) | 481473 (1.9%) | 433291 (2.08%) | 495271 (1.98%) | 479623 (1.95%) |
| **Uniquely mapped** | 21388259 (96.43%) | 25213258 (96.23%) | 24480437 (96.49%) | 19363621 (92.75%) | 23954347 (95.74%) | 23637579 (96.05%) |
| **Read-1** | 10701153 (48.24%) | 12618727 (48.16%) | 12252152 (48.29%) | 9688575 (46.41%) | 11988990 (47.92%) | 11827116 (48.06%) |
| **Read-2** | 10687106 (48.18%) | 12594531 (48.07%) | 12228285 (48.2%) | 9675046 (46.34%) | 11965357 (47.82%) | 11810463 (47.99%) |
| **Reads map to '+'** | 10693846 (48.21%) | 12605858 (48.11%) | 12239256 (48.24%) | 9681823 (46.37%) | 11976675 (47.87%) | 11817990 (48.02%) |
| **Reads map to '-'** | 10694413 (48.21%) | 12607400 (48.12%) | 12241181 (48.25%) | 9681798 (46.37%) | 11977672 (47.87%) | 11819589 (48.03%) |
| **Reads mapped in proper pairs** | 20116998 (90.69%) | 23937836 (91.36%) | 23052036 (90.86%) | 18264546 (87.48%) | 22627440 (90.43%) | 22435994 (91.16%) |
| **Proper-paired reads map to different chrom** | 0 (0%) | 0 (0%) | 0 (0%) | 0 (0%) | 0 (0%) | 0 (0%) |

**Table S45: RT-qPCR primer information**

| Gene | Forward | Reverse | Calculated Efficiency | Reference |
|---|---|---|---|---|
| *16S* | CTGTGCACATCTTGACGGTA | TCAGCGTCAGTTACAGACCA | 81% | Yarwood et al.(1) |
| *hla* | ATGAATCCTGTCGCTAATGCCG | TGACCAGCAATGGTACCTTTCG | 114.04% | Balogh et al. (2) |
| *hlgA* | AGCAGTTGGTTTAATCGCCCCTTT | TTGATGATTTCTGCACCTTGGCCG | 83.86% | Viering et al.(3) |
| *hlgC* | GGTGGTAATTTCCAATCAGCC | GAATGAATTCGCTTTGACGCCC | 109.18% | Balogh et al. (2) |
| *hld/RNAIII* | TAATTAAGGAAGGAGTGATTTCAATG | TTTTTAGTGAATTTGTTCACTGTGTC | 118.36% | Zheng et al.(4) |
| *agrA* | GTGAATTCGTAAGCATGACCCAGTTG | TGTAAGCGTGTATGTGCAGTTTCTAAA | 92.80% | This work |

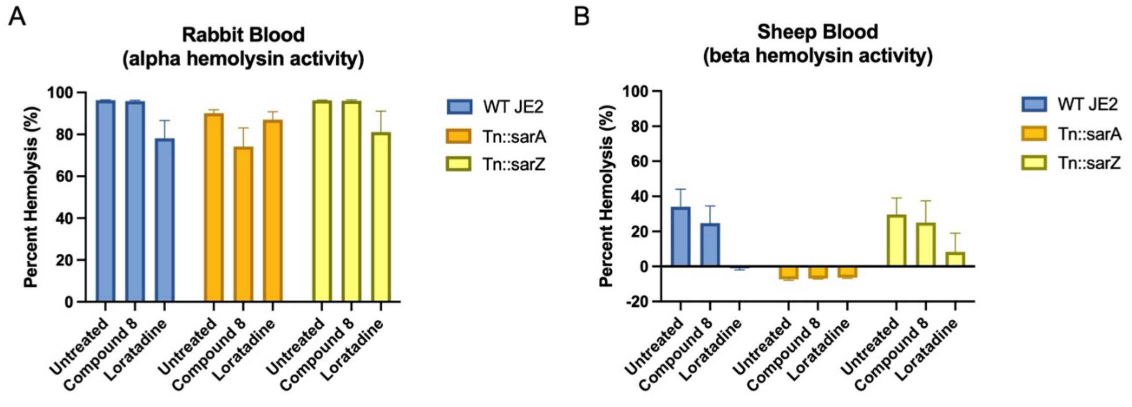

**Figure S1:** Quantitative hemolysis assays using (A) rabbit or (B) sheep blood reveal SarA impacting loratadine-induced reduction of beta hemolytic activity in USA300 strain JE2. Compound 8 results are also included. Mean percent hemolysis is displayed from three biological replicates. Error bars represent the standard error of the mean. Two-way ANOVAs were used to assess variance between means of every sample. Details of this analysis are shared in Tables S1 for loratadine and S6 for compound 8.

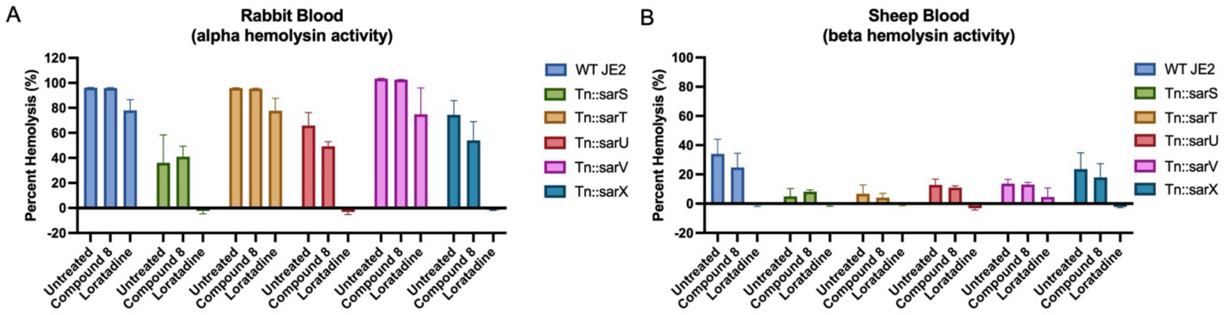

**Figure S2:** Quantitative hemolysis assays using (A) rabbit or (B) sheep blood reveal SarU and SarX inhibiting loratadine's reduction of alpha hemolytic activity in USA300 strain JE2. SarS and SarT impact loratadine's reduction of beta hemolytic activity. Compound 8 results are also included. In both panels, mean percent hemolysis is displayed from three biological replicates. Error bars represent the standard error of the mean. Two-way ANOVAs were used to assess variance between means of every sample. Details of this analysis are shared in Tables S1 for loratadine and S6 for compound 8.

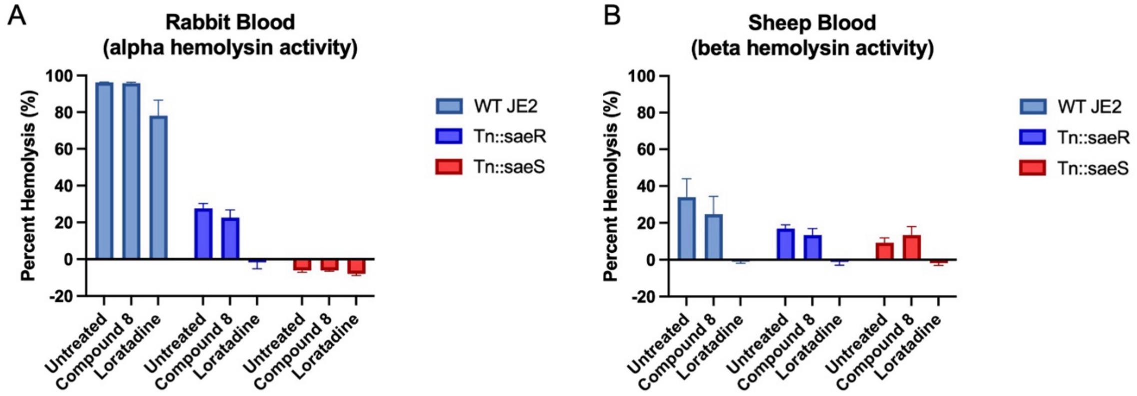

**Figure S3:** Quantitative hemolysis assays using (A) rabbit or (B) sheep blood reveal SaeRS is not required for loratadine's reduction in alpha or beta hemolytic activity in USA300 strain JE2. Compound 8 results are also included. In both panels, mean percent hemolysis is displayed from three biological replicates. Error bars represent the standard error of the mean. Two-way ANOVAs were used to assess variance between means of every sample. Details of this analysis are shared in Tables S1 for loratadine and S6 for compound 8.

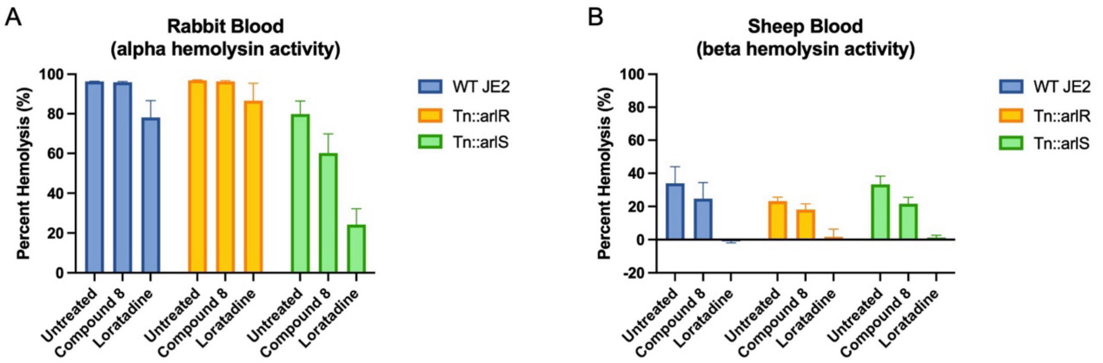

**Figure S4:** Quantitative hemolysis assays using (A) rabbit or (B) sheep blood reveal ArlS inhibiting loratadine's reduction of alpha hemolytic activity in USA300 strain JE2. Compound 8 results are also included. In both panels, mean percent hemolysis is displayed from three biological replicates. Error bars represent the standard error of the mean. Two-way ANOVAs were used to assess variance between means of every sample. Details of this analysis are shared in Tables S1 for loratadine and S6 for compound 8.

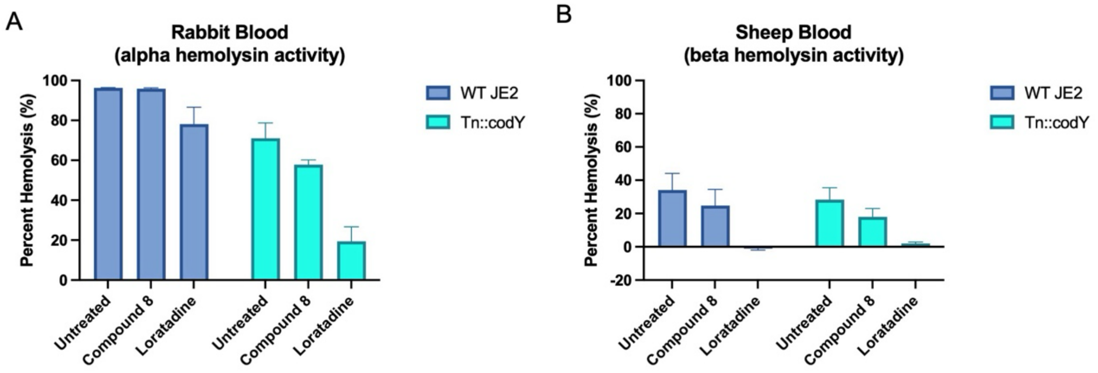

**Figure S5:** Quantitative hemolysis assays using (A) rabbit or (B) sheep blood reveal CodY inhibiting loratadine's reduction of alpha hemolytic activity in USA300 strain JE2. Compound 8 results are also included. In both panels, mean percent hemolysis is displayed from three biological replicates. Error bars represent the standard error of the mean. Two-way ANOVAs were used to assess variance between means of every sample. Details of this analysis are shared in Tables S1 for loratadine and S6 for compound 8.

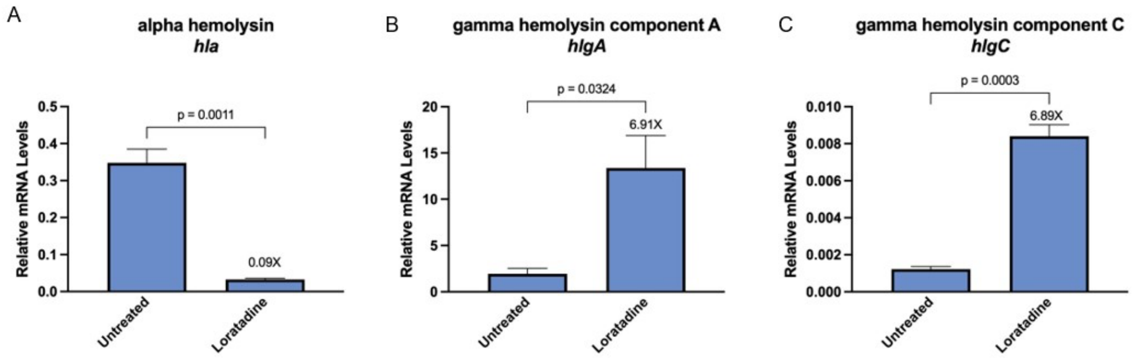

**Figure S6:** RT-qPCR validates multiple hemolysin genes changing with loratadine treatment. (A) *hla* (B) *hlgA* (C) *hlgC*. In all panels, mRNA levels are relative to a 16S reference gene. Error bars represent the standard error of the mean. Labels over treated samples indicate fold change from untreated controls. Unpaired student's t-tests were used to assess statistically significant differences. Resulting p values ≤0.05 are displayed.

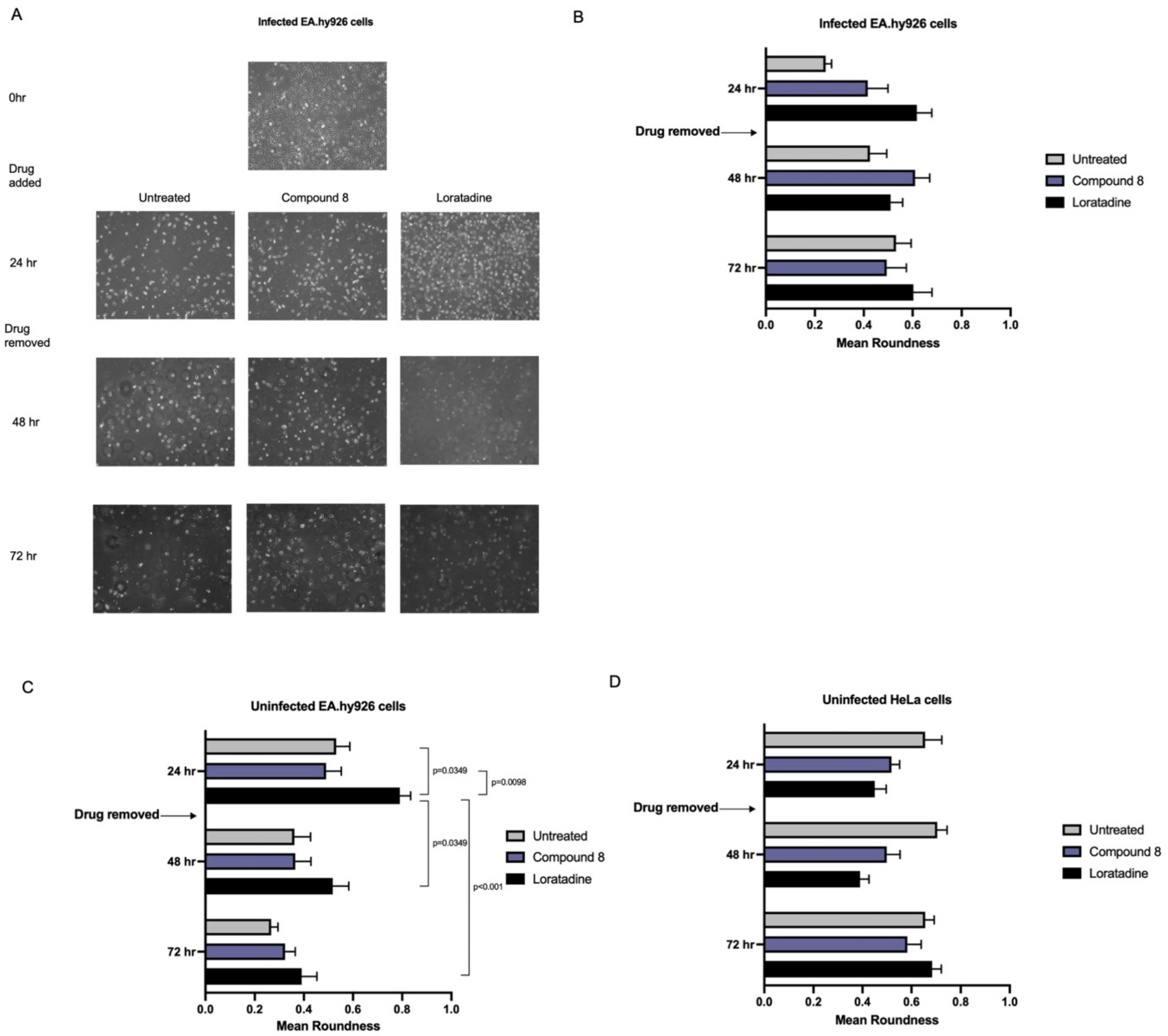

**Figure S7:** Loratadine-induced rounding of endothelial-like EA.hy926 cells is temporary. (A) Brightfield microscopy images of infected EA.hy926 cells over time. (B) Measurement of mean roundness of infected cells shown in panel (A). (C) Measurement of mean roundness of uninfected EA.hy926 cells. (D) Measurement of mean roundness of uninfected HeLa cells (epithelial). In all panels, mean roundness values can range from 0.0-1.0, with 1.0 being a perfect circle. Error bars represent the standard error of the mean. Adjusted p values ≤0.05 are displayed. Details of 2-way ANOVAs are found in Tables S38-40.

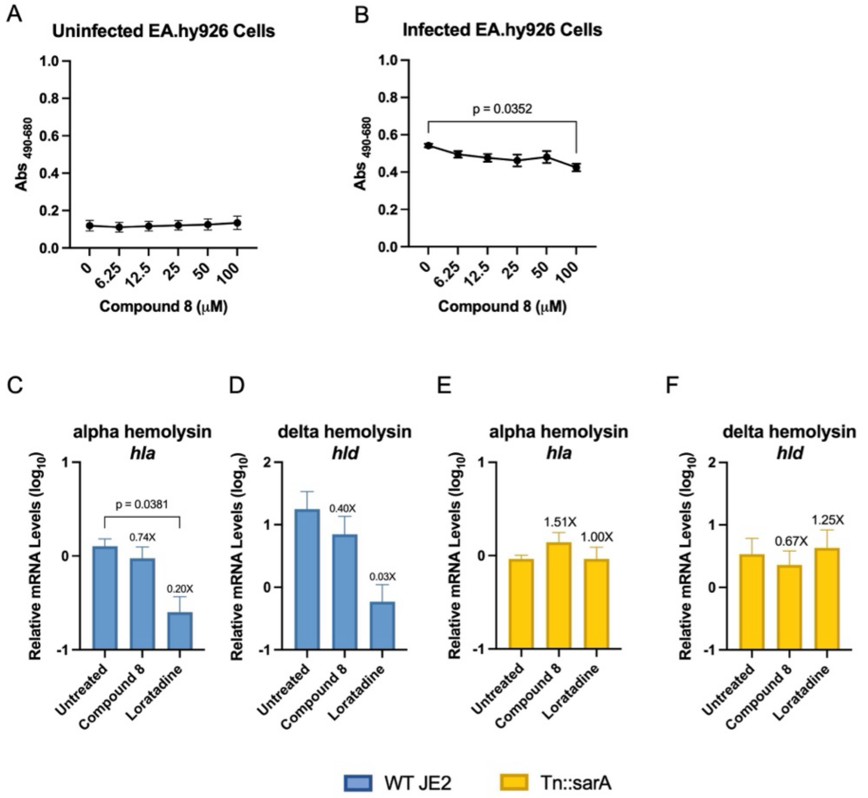

**Figure S8:** Human cell line infection model shows compound 8 does not induce cytotoxicity in a human cell line and multiple hemolysins' mRNA levels get suppressed. (A) Cytotoxicity results of compound 8 in EA.hy926 cells after 24 hrs (B) The same experiment as shown in (A), but in the presence of USA300 JE2 MRSA. Error bars represent the standard error of the mean. RT-qPCR results of (C) *hla* and (D) *hld* levels in internalized USA300 JE2 after 24 hrs of compound 8 treatment. RT-qPCR results of (E*) hla* and (F) *hld* in internalized USA300 JE2 Tn::sarA after 24 hrs of compound 8 treatment. In panels C-F, mRNA levels are relative to a 16S reference gene. Error bars represent the standard error of the mean. Labels over treated samples indicate fold change from untreated controls. Unpaired student's t-tests were used to assess statistically significant differences. Resulting p values ≤0.05 are displayed.

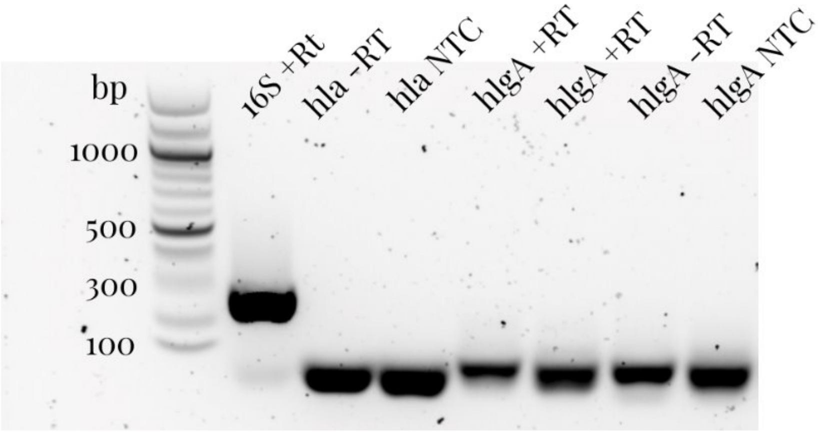

**Figure S9:** Agarose gel electrophoresis of RT-qPCR products. The presence of primer dimers only in hemolysin PCR reactions are verified for internalized USA300 JE2 Tn::stp1. A product of the expected size in 16S reference gene reactions is also verified. Experimental reactions that contained reverse transcriptase are labelled +RT, the controls for genomic DNA contamination are labeled –RT, and those that lacked template are labelled NTC for no template control.
